# Therapeutic suppression of proteolipid protein rescues Pelizaeus-Merzbacher Disease in mice

**DOI:** 10.1101/508192

**Authors:** Matthew S. Elitt, Lilianne Barbar, H. Elizabeth Shick, Berit E. Powers, Yuka Maeno-Hikichi, Mayur Madhavan, Kevin C. Allan, Baraa S. Nawash, Zachary S. Nevin, Hannah E. Olsen, Midori Hitomi, David F. LePage, Weihong Jiang, Ronald A. Conlon, Frank Rigo, Paul J. Tesar

## Abstract

Mutations in *proteolipid protein 1* (*PLP1*) result in failure of myelination and severe neurological dysfunction in the X-linked pediatric leukodystrophy Pelizaeus-Merzbacher disease (PMD). The majority of *PLP1* variants, including supernumerary copies and various point mutations, lead to early mortality. However, *PLP1*-null patients and mice display comparatively mild phenotypes, suggesting that reduction of aberrant *PLP1* expression might provide a therapeutic strategy across PMD genotypes. Here we show, CRISPR-Cas9 mediated germline knockdown of *Plp1* in the severe *jimpy* (*Plp1*^jp^) point mutation mouse model of PMD rescued myelinating oligodendrocytes, nerve conduction velocity, motor function, and lifespan to wild-type levels, thereby validating *PLP1* suppression as a therapeutic approach. To evaluate the therapeutic potential of *Plp1* suppression in postnatal PMD mice, we tested antisense oligonucleotides (ASOs) that stably decrease mouse *Plp1* mRNA and protein *in vivo*. Administration of a single intraventricular dose of *Plp1*-targeted ASOs to postnatal *jimpy* mice increased myelination, improved motor behavior, and extended lifespan through an 8-month endpoint. Collectively, these results support the development of *PLP1* suppression as a disease-modifying therapy for most PMD patients. More broadly, we demonstrate that RNA therapeutics can be delivered to oligodendrocytes *in vivo* to modulate neurological function and lifespan, opening a new treatment modality for myelin disorders.

## Main text

Pelizaeus-Merzbacher disease (PMD; OMIM 312080) is an X-linked leukodystrophy typified by extensive hypomyelination of the central nervous system (CNS). Symptoms typically present early in childhood with a constellation of nystagmus, spasticity, hypotonia, and cognitive dysfunction, leading to mortality in early adulthood. In severe forms, symptoms present connatally and patients succumb to their disease in early childhood. Despite intense interest in PMD therapeutic development, including several clinical trials, no disease modifying therapies have proven efficacious in patients ^1–6^.

Mutations in the *proteolipid protein 1* (*PLP1;* OMIM 300401) gene underlie the pathogenesis of PMD, with variability in disease onset and progression dictated by the severity of the particular mutation ^7–9^. *PLP1* codes for the tetraspan protein PLP as well as its shorter splice isoform DM20, which lacks 35 amino acids in the cytosolic loop region ^10^. PLP/DM20 is highly conserved across species, with human, mouse, and rat sharing identical amino acid sequence ^11^. Expression of *PLP1* is largely restricted to myelinating oligodendrocytes, where it is responsible for ~50% of the total protein content of myelin ^12^.

The majority of PMD cases result from duplications in *PLP1* yielding overexpression of otherwise normal PLP protein *^8^*. Additionally, hundreds of unique point mutations in *PLP1* which each generate abnormal PLP protein have also been discovered in patients with severe disease. Interestingly, while *PLP1* deletions are uncommon, null patients and mice display symptoms that are significantly delayed and more mild compared to those with duplications or point mutations ^13–15^. Null patients can live until 40-60 years old ^8, 16^, do not develop lower body spastic paraparesis until the second or third decade of life, and do not demonstrate cognitive regression until the third or fourth decade of life (see Extended Data Table 1 for detailed clinical presentations of *PLP1*-null patients from published reports).

Collectively, this clinical landscape provides several potential opportunities for therapeutic development. While never tested, knockout or suppression of the duplicated copy of *PLP1* should, in theory, correct elevated protein expression to normal levels in patients harboring duplications. More globally, we suggest that the mild presentation of null patients provides rationale for a generalized *PLP1* suppression strategy that may provide clinical benefit beyond this single use case. Such an approach would provide comprehensive therapy to include the litany of point mutations that generate abnormal PLP protein, abrogating the need for personalized therapy tailored to each patient’s severe, individual mutation.

To test this latter strategy, we evaluated *Plp1* suppression as a therapeutic approach for PMD using the *jimpy* (*Plp1^jp^*) mouse, which expresses a mutant PLP protein due to a point mutation in splice acceptor site of intron 4 of *Plp1*, leading to exon 5 skipping and a frameshift affecting the final 70 amino acids in the C-terminus of PLP/DM20. *Jimpy* mice exhibit severe neurological symptoms, die at 3 weeks of age, and accurately reflect the cellular and molecular pathology seen in human disease. To test if *Plp1* suppression was a valid therapeutic strategy for PMD we first utilized nuclease-mediated cleavage of DNA by clustered regularly interspaced short palindromic repeat-associated 9 (CRISPR-Cas9) protein ^17,18^ with synthetic guide RNAs (sgRNA) targeted to induce a frameshift in PLP1 exon 3 and trigger subsequent knockdown of *Plp1* mRNA by nonsense-mediated decay. *Plp1*-targeting sgRNAs were designed and tested for on-target cutting efficiency and two sgRNAs (3 and 7) were administered to knockdown *Plp1* by germline editing of *jimpy* mice (Fig. 1a). Concomitant administration of both sgRNAs with Cas9 mRNA from Streptococcus pyogenes (spCas9) in zygotes from crosses of *jimpy*-carrier females with wild-type males generated mice with high on-target mutation efficiency in *Plp1* (Extended Data Fig. 1a). These founders included a *jimpy* male with complex deletion of 80 total nucleotides of DNA in exon 3 of his single copy of *Plp1* on the X chromosome, resulting in a frameshift in exon 3 and an early termination codon in exon 4 (Fig. 1a and Extended Data Fig. 1b). This mouse denoted “CR-*impy*” (for <u>CR</u>ISPR frameshift-mediated knockdown of *Plp1* in *j<u>impy</u>*) showed no overt neurological phenotypes. A line of CR-*impy* was bred to evaluate cellular, molecular, and functional phenotypes, along with contemporaneous, isogenic wild-type and *jimpy* male mice.

**Fig. 1.**
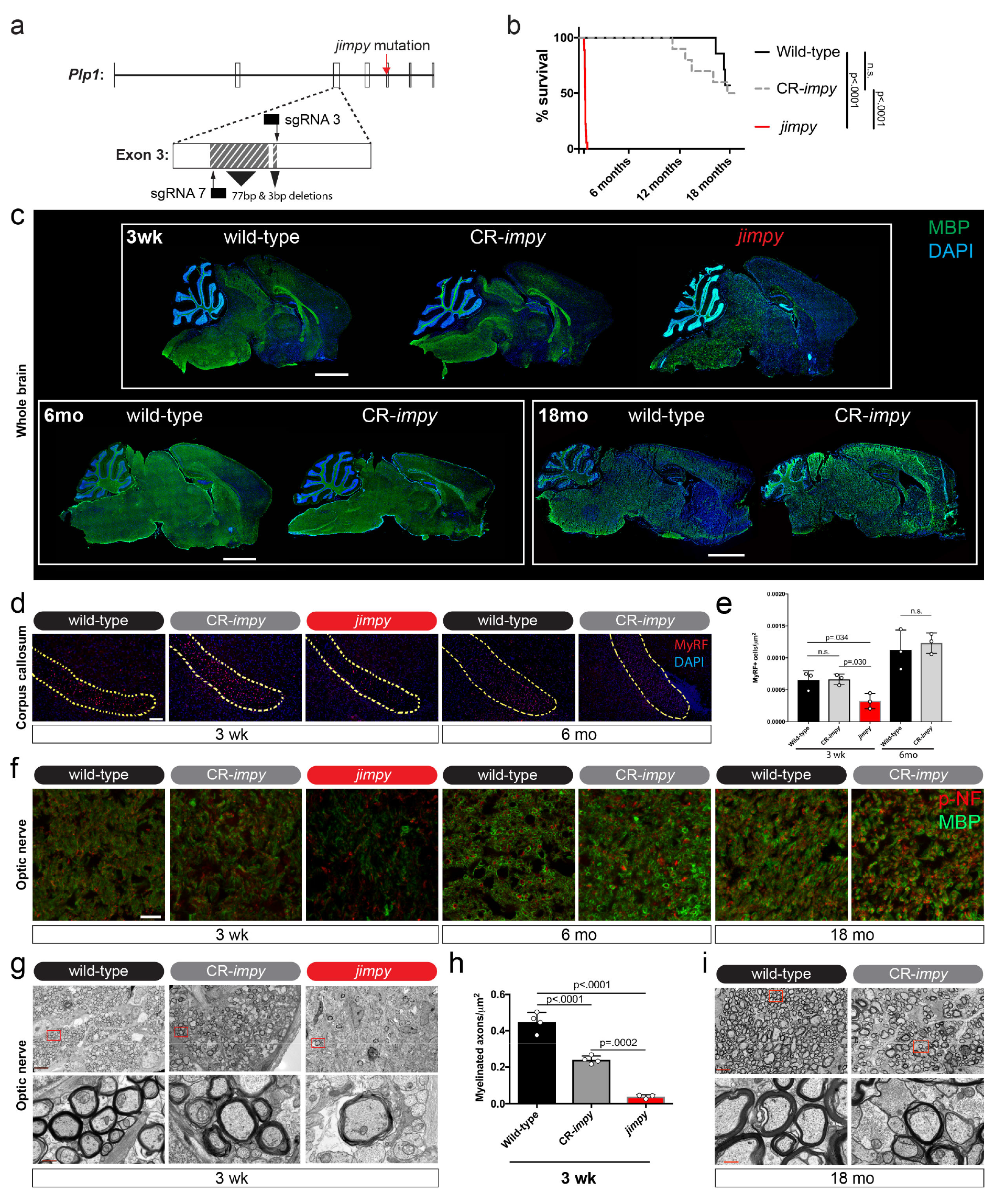
Prevention of PMD and rescue of lifespan by CRISPR-mediated knockdown of *Plp1* in the germline of *jimpy* mice. **a**, Schematic of *Plp1* targeting in *jimpy* using CRISPR-spCas9. The location of the *jimpy* mutation in the 3’ splice acceptor site of intron 4 is indicated by a red arrow. Two separate sgRNAs (denoted 3 and 7) were utilized and their DNA binding sites are indicated by solid black bars and their predicted cutting sites by black arrows. The complex, frameshift deletion of 80 base-pairs of deleted sequence within exon 3 in the *CR-*impy** mice is shown by two grey hashed boxes (see Extended Data Fig. 1 for more detail and sequence traces). **b**, Kaplan-Meier plot comparing lifespan of contemporaneous wild-type, *jimpy*, and CR-*impy* mouse cohorts. n = 25, 18, and 23 starting animals in the wild-type, *jimpy*, and CR-*impy* cohorts, respectively (see Supplementary Fig. 1 for metadata of every animal in this study including censoring of animals for molecular and functional studies at pre-determined time points). p-values calculated using the log-rank test between cohorts. **c**, Representative immunohistochemical images of whole-brain sagittal sections showing MBP+ oligodendrocytes (green) and total DAPI+ cells (blue) in wild-type, CR-*impy*, and *jimpy* mice at 3 weeks, 6 months, and 18 months of age. Scale bar, 2mm. **d**, Representative sagittal immunohistochemical images of the rostral end of the corpus collosum showing MyRF+ oligodendrocytes (red) and total DAPI+ cells (blue) in wild-type, CR-*impy*, and *jimpy* mice at 3 weeks, 6 months, and 18 months of age. Scale bar, 100μm. **e**, Quantification of MyRF+ oligodendrocytes in the corpus callosum at 3 weeks and 6 months of age for each genotype. n=3 mice per genotype at each time point. Error bars show mean ± standard deviation. p-values calculated using ANOVA with Tukey correction for multiple comparisons at 3 weeks and Welch’s t-test at 6 months. **f**, Representative confocal immunohistochemical images of optic nerve cross sections showing MBP+ oligodendrocytes (green) and phospho-neurofilament+ intact axons (red) in wild-type, CRimpy, and *jimpy* mice at 3 weeks, 6 months, and 18 months of age. Scale bar, 20μm. **g**, Representative electron micrograph images showing myelination in optic nerve cross sections in wild-type, CR-*impy*, and *jimpy* mice at 3 weeks of age. Lower panel is a higher magnification of red boxed area in the upper panel. Upper panel scale bar, 5μm and lower panel scale bar is 0.5μm. **h**, Quantification of the number of myelinated axons in optic nerves of each genotype at postnatal week 3. n=4 mice for wild-type and CR-*impy* and n=3 mice for and *jimpy* genotypes. Error bars show mean ± standard deviation. p-values calculated using ANOVA with Tukey correction for multiple comparisons. **i**, Representative electron micrograph images of myelination in optic nerve cross sections in wild-type and CR-*impy* mice at 18 months of age. Lower panel is a higher magnification of red boxed area in the upper panel. Upper panel scale bar, 5μm and lower panel scale bar is 0.5μm.

As expected *jimpy* males displayed the typical pathological phenotypes including severe tremor, ataxia, seizures (lasting >30 seconds), and death by the third postnatal week (Fig. 1b, Supplementary Fig. 1, Supplementary Video 1). In contrast, CR-*impy* mice, which had a 61-74% reduction *Plp1* transcript expression in multiple brain regions (Extended Data Fig. 1c), showed a 21-fold increase in lifespan (mean survival = 23 and 489 days in *jimpy* and CR-*impy*, respectively) with no overt tremor, ataxia, or evidence of seizures through the terminal endpoint at 18 months of age (Fig. 1b, Supplementary Fig. 1, Supplementary Video 1, and Supplementary Video 2).

Assessment of CR-*impy* mice at 3 weeks, 6 months, and 18 months of age by immunohistochemistry revealed a stable restoration of oligodendrocytes to near wild-type levels throughout the neuraxis, as evidenced by expression of the mature myelin marker myelin basic protein (MBP), and quantification of oligodendrocyte number using the oligodendrocyte-specific transcription factor myelin regulatory factor (MyRF) (Fig. 1c-e). Quantification of MBP protein by western blot and RNA by qRT-PCR further supported these findings, revealing 40-95% restoration of MBP at 3 weeks and full restoration of *Mbp* expression and MBP at 6 months of age across multiple brain regions in CR-*impy* mice compared to wild-type (Extended Data Fig. 1d-f). Restoration of oligodendrocytes in CR-*impy* mice grossly reduced reactivity of microglia and astrocytes normally seen in *jimpy* mice at 3 weeks of age, and was sustained into adulthood (Extended Data Fig. 2a, b). Finally, MBP+ CR-*impy* oligodendrocytes ensheathed phospho-neurofilament+ axons in a similar manner to wild-type samples (Fig. 1f), suggestive of a potentially comparable myelination status.

Electron micrographs and tissue sections stained with toluidine blue demonstrated increased myelination in *CR-*impy** mice. Examination of optic nerves, which provide straightforward quantification of myelination from aligned axons, revealed a significant 6-fold improvement in the number and density of myelinated axons in *CR-*impy** animals, reaching 53% of wild-type levels at 3 weeks of age (Fig. 1g, h, and Extended Data Fig. 3a). Myelination in CR-*impy* animals was sustained through 6 month and 18 month time points, approaching wild-type levels (Fig. 1i and Extended Data Fig. 3b, c). Additionally, examination of corpus callosum myelin at 3 weeks (Extended Data Fig. 3d) further revealed that these improvements were not restricted to the optic nerve but manifested throughout the neuraxis of CR-*impy* mice.

Although the quantity of myelin increased drastically compared to *jimpy*, we noted that CR*-impy* myelin sheaths were not quite as compact as those seen in age-matched wild type controls. To determine if restored myelin in CR-*impy* mice was functional we used electrophysiology to measure conduction velocity in the optic nerve at two time points. At 3 weeks of age we found a significant 2.6- and 1.7-fold increase in 1^st^ and 2^nd^ peak conduction velocities, respectively, in CR-*impy* mice compared to *jimpy*. Intriguingly, these values trailed wild-type conduction velocities, a finding consistent with our quantitative electron microscopy showing an incomplete restoration of myelinated axons at the 3 week time point. However, over time this discrepancy disappeared and by 6 months of age we found no significant difference in optic nerve conduction velocity in CR-*impy* mice relative to wild-type (Fig. 2a, b), suggesting that suppression of *Plp1* does not impair myelin function.

**Fig. 2.**
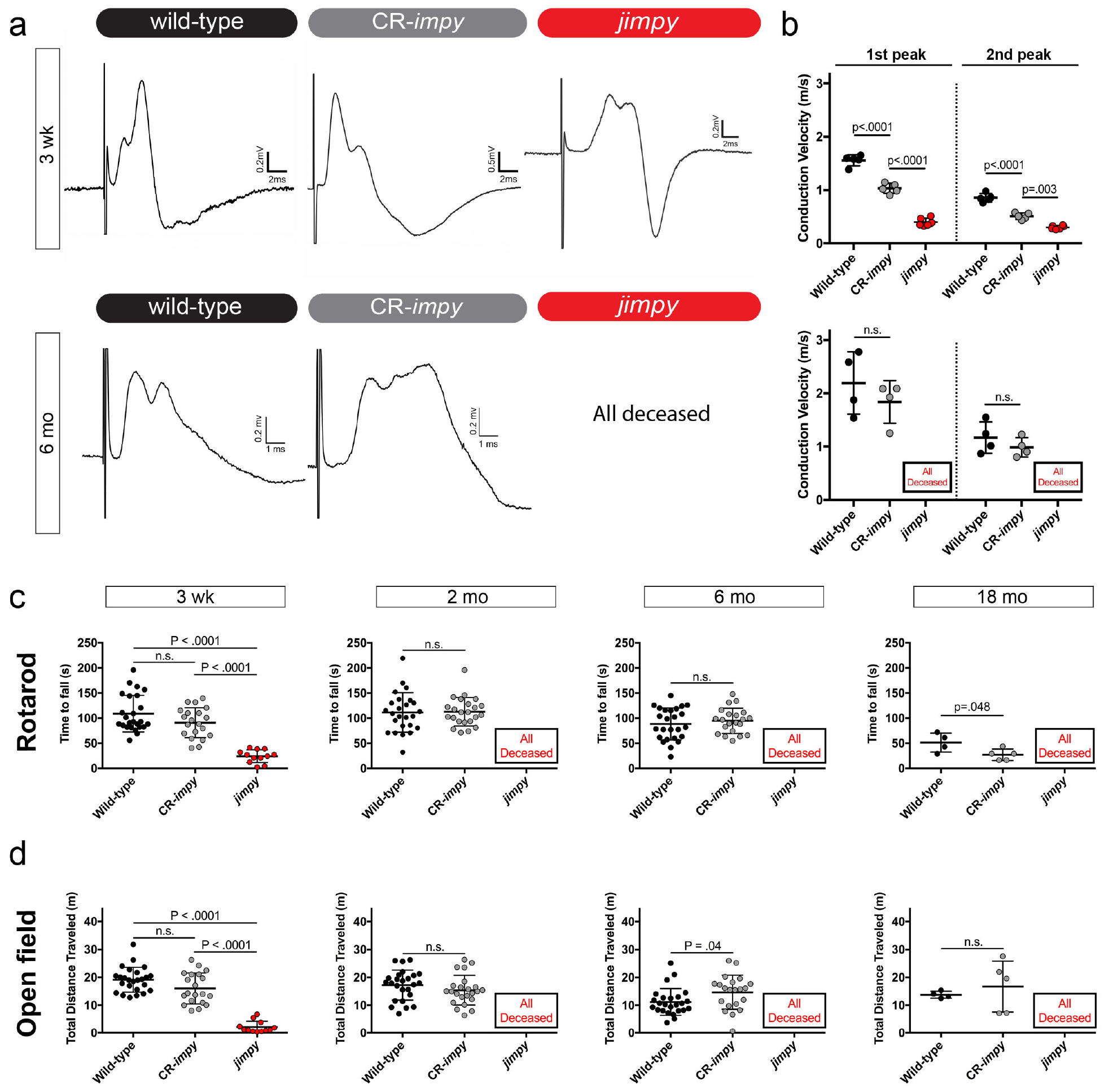
CRISPR-mediated knockdown of *Plp1* in *jimpy* mice restores myelin function and motor phenotypes. **a**, Representative electrophysiology optic nerve conduction traces from wild-type, *CR-*impy*, jimpy* mice at 3 weeks and 6 months of age. **b**, Quantification of optic nerve conduction velocities from wild-type, CR-*impy, jimpy* mice at 3 weeks and 6 months of age. Each point represents an individual biological replicate (optic nerves from separate mice) with n = 5, 5, and 6 wild-type, *jimpy*, and CR-*impy* mice, respectively, at the 3 week time point and n = 4 wild-type and CR-*impy* mice each at the 6 month time point. Error bars show mean ± standard deviation. p-values calculated using one-way ANOVA with Tukey correction for multiple comparisons at 3 weeks and two-way, unpaired t-test at 6 months. **c**, Comparison of motor function and coordination of wild-type, CR-*impy*, and *jimpy* mice at 3 weeks (n=25, 20, and 12 wild-type, *jimpy*, and CR-*impy* mice), 2 months (n= 25 and 23 wild-type and CR-*impy* mice), 6 months (n=25 and 21 wild-type and CR-*impy* mice), and 18 months of age (n=4 and 5 wild-type and CR-*impy* mice) by accelerating rotarod performance. Individual data points represent the mean time to fall of three separate trials for each biological replicate (separate mice). Error bars show mean ± standard deviation. p-values calculated using one-way ANOVA with Tukey correction for multiple comparisons at 3 weeks or two-way, unpaired t-test at later time points. See Supplementary Fig. 1 for raw data values. **d**, Comparison of locomotor activity of wild-type, CR-*impy*, and *jimpy* mice at 3 weeks (n=25, 20, and 12 wild-type, *jimpy*, and CR-*impy* mice), 2 months (n= 25 and 23 wild-type and CR-*impy* mice), 6 months (n=25 and 21 wild-type and CR-*impy* mice), and 18 months of age (n=4 and 5 wild-type and CR-*impy* mice) by open field testing. Individual data points represent total distance traveled for each biological replicate (separate mice). Error bars show mean ± standard deviation. p-values calculated using one-way ANOVA with Tukey correction for multiple comparisons at 3 weeks or two-way, unpaired t-test at later time points. See Supplementary Fig. 1 for raw data values.

Finally, we wanted to determine if the widespread restoration of CR-*impy* oligodendrocytes resulted in a meaningful recovery of motor performance using the accelerating rotarod and open field behavioral assays at 3 weeks, 2 months, 6 months, and 18 months of age. In rotarod testing, which measures motor learning, function, and coordination, CR*-impy* mice showed equivalent performance to wild-type up through 6 months of age. At the final 18 month time point we noted a slight decrease in CR-*impy* performance relative to wild-type (Fig. 2c). This is consistent with prior reports in mice with complete *Plp1* knockout in otherwise wild-type mice, which showed a decline in rotarod performance in late adulthood relative to wild-type due to long tract axon degeneration ^13^. Overall locomotion of CR-*impy* mice in open field testing was equivalent to wild-type at all time points tested (Fig. 2d). Together, these results establish that frameshift-mediated knockdown of mutant *Plp1* in PMD mice prevents disease with near complete restoration of oligodendrocytes, functional myelin, and lifespan.

PMD is thought to result from a cell-intrinsic deficit within the oligodendrocyte lineage ^19^-^23^. In the CNS, PLP (protein) is restricted to oligodendrocytes, but *Plp1* transcript and transgene expression have been reported in a few neuronal subsets ^24^ Since CR-*impy* mice have constitutive germline *Plp1* knockdown in all cells, we generated and validated induced pluripotent stem cells (iPSCs) from isogenic wild-type, CR-*impy*, and *jimpy* mice (Extended Data Fig. 4a, b) to generate pure populations of oligodendrocyte progenitor cells (OPCs) and assess the cell type specific effect of *Plp1* knockdown, *in vitro* ^25,26^. OPCs expressing the canonical transcription factors Olig2 and Sox10 (Extended Data Fig. 4c, d) were stimulated to differentiate towards an oligodendrocyte fate by addition of thyroid hormone, and MBP+ oligodendrocytes were quantified. As expected ^22^, *jimpy* cultures contained only rare surviving MBP+ oligodendrocytes with a concomitant loss in total cells (Fig. 3a-c). In contrast CR-*impy* cultures showed a complete rescue of MBP+ oligodendrocytes as well as total cells (Fig. 3a-c). These results suggest that oligodendrocyte restoration in CR*-impy* mice is due to an amelioration of intrinsic cellular pathology within the oligodendrocyte lineage.

**Fig. 3.**
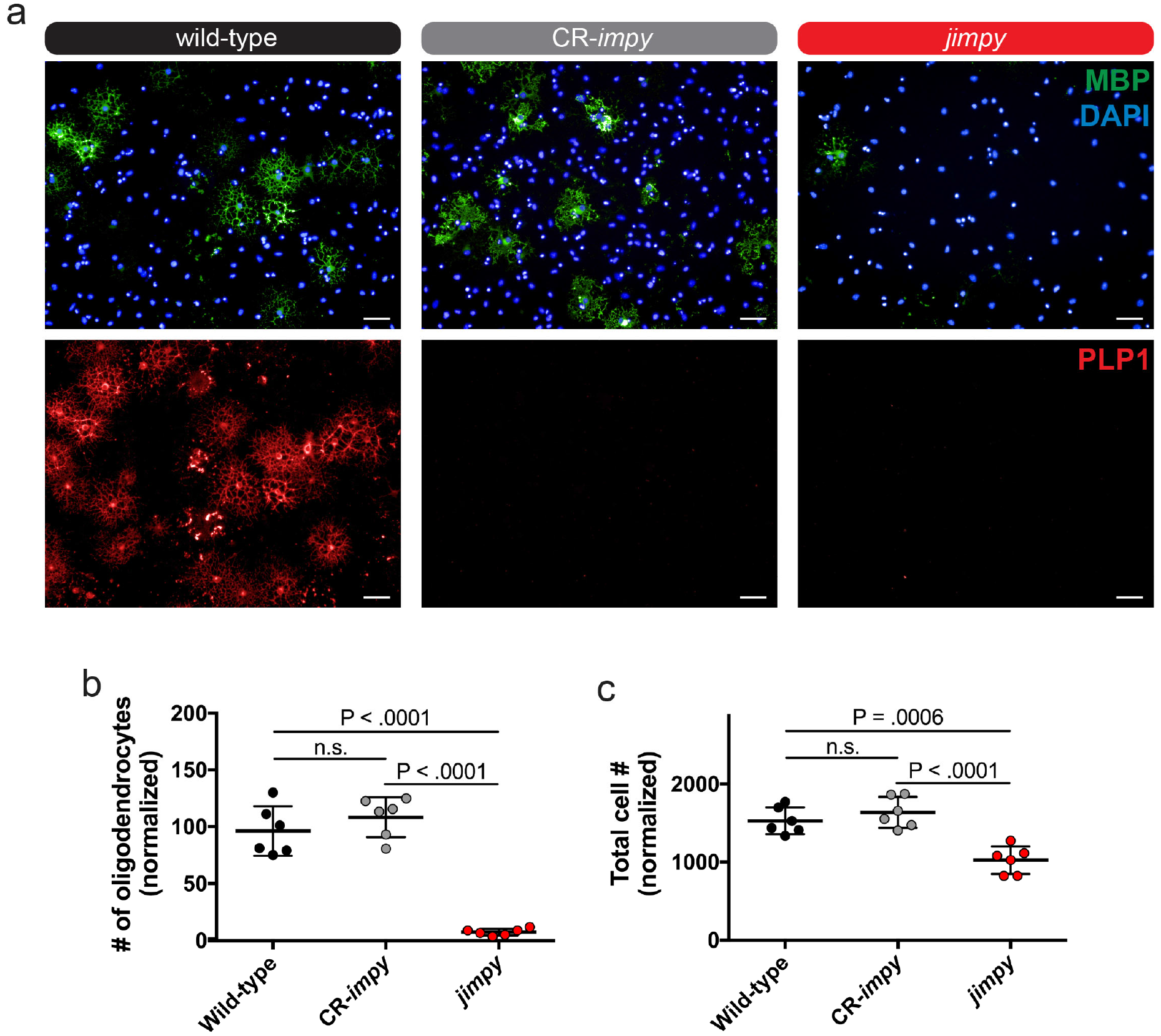
CRISPR-mediated knockdown of *Plp1* in *jimpy* OPCs rescues survival of differentiating oligodendrocytes *in vitro*. **a**, Representative immunocytochemistry images of MBP+ oligodendrocytes and PLP+ oligodendrocytes from wild-type, CR-*impy*, and *jimpy* iPSC-derived OPCs differentiated *in vitro* for 3 days. Top and bottom rows are the same field but channels are separated for clarity. Note the PLP1 antibody detects a C-terminal peptide sequence not present in jimpy and therefore absence of staining simply serves as validation of jimpy genotype but not quantification of total PLP1 protein (for which there are no validated N-terminal antibodies for immunostaining). Scale bar, 50μm. **b-c**, Quantification of MBP+ oligodendrocytes (b) and total DAPI+ cells (c). Error bars show mean ± standard deviation. n=6 technical replicates (single cell line per genotype with 6 separate wells scored). p-values calculated using one-way ANOVA with Tukey correction for multiple comparisons.

**Fig. 4.**
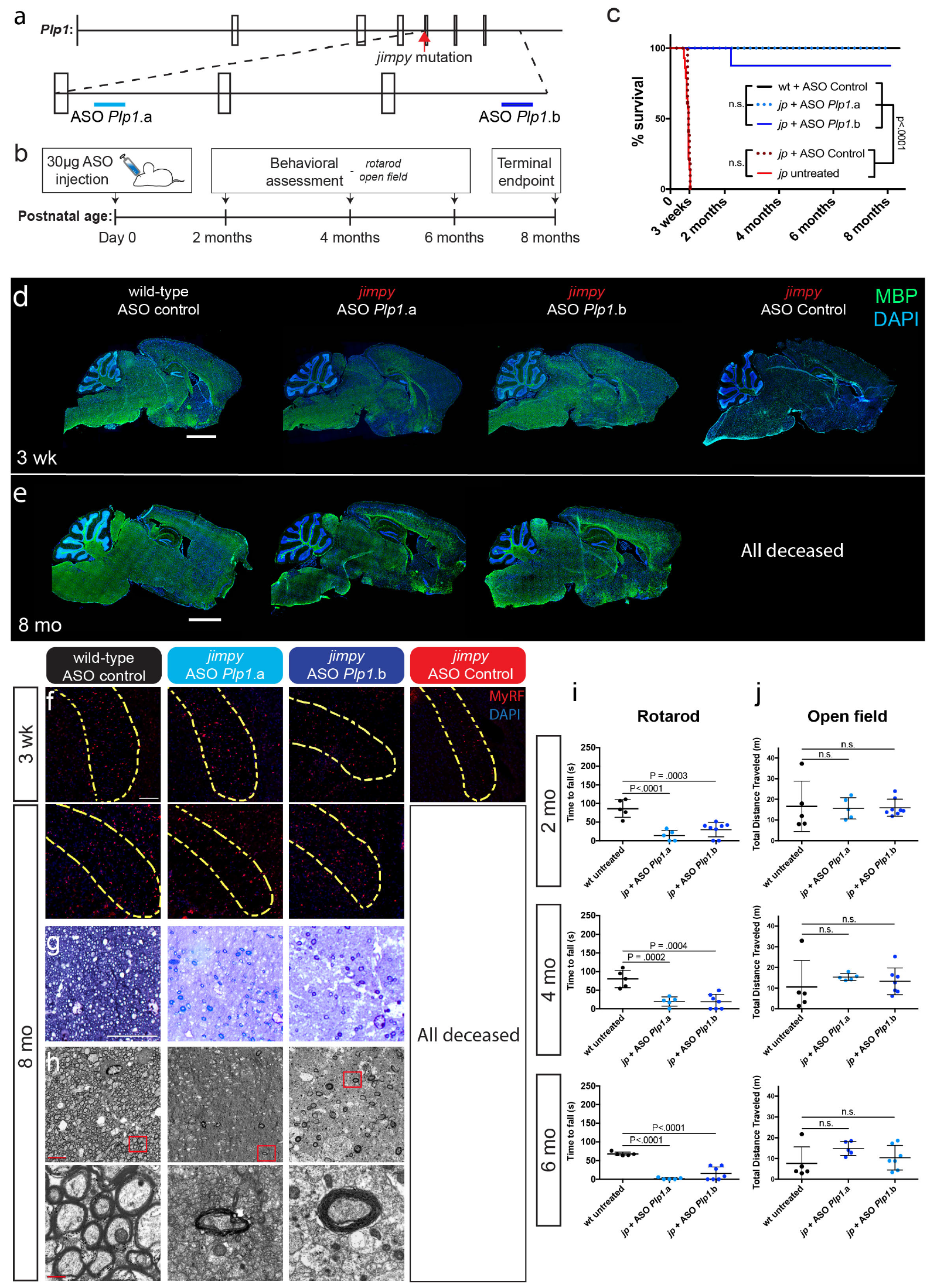
Postnatal delivery of *Plp1*-targeted antisense oligonucleotides rescues lifespan and partially restores functional myelinating oligodendrocytes in *jimpy* mice. **a**, Schematic of the binding location of the two independent ASOs within intron 5 and the 3’ UTR of the *Plp1* pre-mRNA. The location of the *jimpy* mutation in the 3’ splice acceptor site of intron 4 is indicated by a red arrow. **b**, Schematic of the experimental design for ASO experiments. A single 30ug dose of ASO was administered by intracerebroventricular injection into the lateral ventricle within one day after birth. Functional assessment was performed at 2, 4, and 6 months of age and the experiment was terminated for histological analyses at 8 months of age. **c**, Kaplan-Meier plot depicting the survival of *jimpy* mice treated with two independent *Plp1*-targeting ASOs compared to controls. Groups included: untreated wild-type (n=5), wild-type treated with ASO-control (n=12), untreated *jimpy* (n=14), *jimpy* treated with ASO-control (n=5), *jimpy* treated with ASO-*Plp1*.a (n=5), and *jimpy* treated with ASO-*Plp1*.b (n=8). See Supplementary Fig. 3 for metadata of every animal in this study including behavioral studies at pre-determined time points. p-values calculated using the log-rank test. **d-e**, Representative immunohistochemical images of 3 week (d) and 8 month (e) whole-brain sagittal sections showing MBP+ oligodendrocytes (green) and total DAPI+ cells (blue) in control and *jimpy* mice treated with indicated ASOs. Scale bar, 2mm. **f**, Representative sagittal images of the 3 week old rostral corpus callosum showing MyRF+ oligodendrocytes in control and *jimpy* mice treated with indicated ASOs. Scale bar, 100um. **g**, Representative toluidine blue stained images showing myelinated axons in the corpus callosum of 8 month old wild-type and *jimpy* mice treated with the indicated ASOs. Scale bar, 20um. **h**, Representative electron micrograph images showing myelination in the corpus callosum of 8 month old wild-type and *jimpy* animals treated with the indicated ASOs. Lower panel is a higher magnification of red boxed area in the corresponding image in the upper panel. Upper panel scale bar, 5um and lower panel scale bar is 0.5um. **i**, Comparison of motor function and coordination of 2 month (n=5, 5, and 8 for untreated wild-type, ASO-*Plp1*.a-treated *jimpy*, and ASO-*Plp1*.b-treated *jimpy* mice), 4 month (n=5, 5, and 7 for untreated wild-type, ASO-*Plp1*.a-treated *jimpy*, and ASO-*Plp1*.b-treated *jimpy* mice), and 6 month (n=5, 5, and 7 for untreated wild-type, ASO-*Plp1*.a-treated *jimpy*, and ASO-*Plp1*.b-treated *jimpy* mice) old mice by accelerating rotarod performance. Individual data points represent the mean time to fall of three separate trials for each biological replicate (separate mice). Error bars show mean ± standard deviation. p-values calculated using one-way ANOVA with Dunnett’s correction for multiple comparisons. See Supplementary Fig. 3 for raw data values. **j**, Comparison of locomotor activity of 2 month (n=5, 5, and 8 for untreated wild-type, ASO-*Plp1*.a-treated *jimpy*, and ASO-*Plp1*.b-treated *jimpy* mice), 4 month (n=5, 5, and 7 for untreated wild-type, ASO-*Plp1*.a-treated *jimpy*, and ASO-*Plp1*.b-treated *jimpy* mice), and 6 month (n=5, 5, and 7 for untreated wild-type, ASO-*Plp1*.a-treated *jimpy*, and ASO-*Plp1*.b-treated *jimpy* mice) old mice by open field testing. Individual data points represent total distance traveled for each biological replicate (separate mice). Error bars show mean ± standard deviation. p-values calculated using one-way ANOVA with Dunnett’s correction for multiple comparisons. See Supplementary Fig. 3 for raw data values.

After genetically validating *Plp1* knockdown as a therapeutic target for disease-modification in PMD, we pursued a clinically translatable strategy for *in vivo*, postnatal suppression of *Plp1*. While postnatal delivery of CRISPR-based therapeutics has demonstrated pre-clinical efficacy in a separate CNS disorder ^27^, delivery challenges, off-target risks ^28,29^, and the potential to generate more severe, in-frame mutations due to imprecise repair ^30,31^ led us to employ anti-sense oligonucleotides (ASOs) to test suppression of *Plp1*. ASOs are short single-stranded oligodeoxynucleotides with chemical modifications that confer enhanced pharmacological properties including robust *in vivo* stability, target affinity, and cellular uptake when delivered directly to the CNS since they do not cross the blood-brain barrier ^32,33^. ASOs bind to their target RNAs through complementary base pairing and can be designed to modify RNA splicing or form an ASO/RNA hybrid that is recognized by RNase H1, leading to cleavage of the target transcript and concomitant reduction in protein expression. Recently, ASOs have shown remarkable efficacy in several animal models of neuron-based CNS disorders and human spinal muscular atrophy patients, the latter leading to the first FDA-approved therapy for this disease ^34^-^44^ Whether ASOs could be delivered to oligodendrocytes *in vivo* and mediate functional improvement in the context of myelin disease was unknown.

We tested two separate RNase H ASOs targeting the 5^th^ intron (ASO Plp1.a) and 3’UTR (ASO *Plp1.b*) of *Plp1* (Fig. 4a). Administration of ASO *Plp1.a* or ASO *Plp1.b* by intracerebroventricular (ICV) injection robustly reduced *Plp1* expression by 93% and 86% in the cortex and 97% and 94% in the spinal cord of adult wild-type mice, respectively (Extended Data Fig. 5a). Both of these *Plp1*-targeting ASOs were well-tolerated based on CNS histology and lack of alteration or reactivity of glial and immune cell markers by qRT-PCR and immunohistochemistry 8 weeks after dosing adult wild-type mice (Extended Data Fig. 5b-f).

Administration of ASO *Plp1*.a, ASO *Plp1*.b, or a non-targeting control ASO by ICV injection to male pups after birth revealed widespread distribution and stability of ASOs throughout the neuraxis in both wild-type and *jimpy* mice based on whole-brain immunohistochemical staining at 3 weeks (Fig. 4a, b, and Extended Data Fig. 6a, b). In wild-type mice, *Plp1*-targeting ASOs delivered with this single dose treatment regimen showed robust reductions in *Plp1* transcript by 46-90% and PLP protein by 47-63% across multiple CNS regions, but importantly had no effect on MBP protein levels or overt phenotype (Extended Data Fig. 7a-c and Supplementary Fig. 3). As expected, *jimpy* mice treated with non-targeting control ASO and those left untreated succumbed to their disease at the third postnatal week (Fig. 4c). However, *jimpy* mice treated after birth with a single ICV dose of *Plp1*-targeting ASOs (*Plp1*.a or *Plp1*.b) induced a remarkable extension of lifespan, to our terminal endpoint of 8 months of age (when all animals were processed for histology) (Fig. 4b, c, Supplementary Fig. 3, Supplementary Video 3, and Supplementary Video 4).

Treatment with *Plp1*-targeting ASOs increased oligodendrocytes in *jimpy* animals by 3 weeks of age, notably in the brainstem, which was sustained through the 8 month terminal end point without additional ASO dosing or other intervention (Fig. 4d-f and Extended Data Fig. 7d, e). Electron micrographs and tissue sections stained with toluidine blue confirmed that some myelinated axons were still present even 8 months post-treatment, but overall myelination was reduced relative to wild-type controls (Fig. 4g, h). Symptomatically, *Plp1*-targeting ASO-treated *jimpy* mice showed only minor PMD pathological phenotypes, including slight tremor and occasional short duration seizures (<15 seconds), but otherwise appeared overtly normal in daily activities including the ability to breed, which has not previously been achieved by a *jimpy* male mouse (Supplementary Fig. 3). Rotarod performance of *Plp1*-targeting ASO-treated *jimpy* mice lagged below wild-type levels, but, strikingly, overall locomotion was restored to wild-type levels across 2 month, 4 month, and 6 month time points (Fig. 4i, j). Together these data demonstrate that a single postnatal administration of ASOs elicits a sustained reduction in *Plp1* expression and dramatically improves myelination, motor performance, and lifespan in a severe point mutation model of PMD.

In summary, we have shown CRISPR- and ASO-mediated rescue of PMD in the severely affected *jimpy* mouse model through two independent, technological modalities to achieve mutant *Plp1* suppression. We demonstrate that RNA-based drugs can be used to modulate a disease target in oligodendrocytes to restore both functional myelin and lifespan in the context of a severe genetic disorder. These results provide powerful foundational data for the development of clinically relevant ASO technology to achieve postnatal reduction of *Plp1*. While further pre-clinical development is needed to optimize the dosing regimen, our results highlight that even a single ASO treatment can elicit a profound and sustained phenotypic improvement.

The genetic spectrum of PMD patients encompasses hundreds of unique mutations. Our data nominates a mutation-agnostic approach to collapse this heterogeneity through suppression of *PLP1*, potentially abrogating the need for per-patient, personalized therapies. Importantly, while mice display minimal phenotype when *Plp1* is knocked out ^13,14,45^, *PLP1*-null patients present with neurological disease, but with considerably later onset, slower progression, and improved clinical outcomes. While careful titration of normal *PLP1* expression to wild-type levels could be curative for the majority of patients who harbor gene duplications, reducing mutant *PLP1* in patients with point mutations, while not a full cure, may provide substantial improvement for this disease, which currently has no viable treatment options. Collectively our studies, combined with the feasibility of ASO delivery to the human CNS and current safety data in other CNS indications, support advancement of *PLP1* suppression into the clinic as a disease modifying therapeutic with potential universal applicability to PMD patients. More broadly, our data provide a framework to modulate pathogenic genes or mRNAs in OPCs and oligodendrocytes in order to restore myelination and neurological function in additional genetic and sporadic disorders of myelin.

## Acknowledgments

This research was supported, in part, by grants from the NIH R01NS093357 (P.J.T.), T32GM007250 (M.S.E, Z.S.N., K.C.A), and F30HD084167 (Z.S.N.); the New York Stem Cell Foundation (P.J.T); the European Leukodystrophy Association (P.J.T.); and philanthropic contributions from the Geller, Goodman, Fakhouri, Long, Peterson, and Weidenthal families. Additional support was provided by the Genomics, Small Molecule Drug Development, and Rodent Behavioral core facilities of the Case Western Reserve University (CWRU) Comprehensive Cancer Center (P30CA043703), the CWRU Light Microscopy Imaging Center (S10-OD016164), and the electron microscopy division of the Cleveland Clinic Lerner Research Institute Imaging Core. We are grateful to Lynn Landmesser, Robert Miller, Peter Scacheri, Tony Wynshaw-Boris, Ben Clayton, Simone Edelheit, Alex Miron, Hiroyuki Arakawa, Lucille Hu, Chris Allan, and Jared Cregg for technical assistance and discussion.

## Author contributions

M.S.E. and P.J.T. conceived and managed the overall study. H.E.S and M.S.E. maintained the animal colonies, tracked survival, and harvested animal tissues. M.S.E. captured video recordings. L.B. and M.S.E. designed and tested sgRNAs. D.F.L., R.A.C., and W.J. performed zygote electroporation and oviduct transfers. H.E.S., B.S.N., K.C.A., and L.B. performed western blotting and protein quantitation. B.E.P, L.B., and K.C.A. performed qRT-PCR. M.M., B.S.N, L.B., H.E.S., and M.S.E. generated the immunohistochemistry data. Y.M. performed optic nerve electrophysiology studies. Y.M., M.H., and H.E.S. processed samples for histology and electron microscopy and analyzed images. M.S.E, K.C.A., B.S.N., and L.B. performed animal behavior. M.S.E., B.S.N., and H.E.O. generated and characterized iPSCs and OPCs in vitro. B.E.P. and F.R. designed and characterized ASOs, tested tolerability in adult mice, recommended the use of ASOs, and contributed to the study design and interpretation of results in the ASO-treated disease model. M.S.E. performed ASO injections in pups. Z.S.N. contributed key components to experimental design, data analysis, and manuscript composition. M.S.E., M.M., L.B., and P.J.T. assembled figures. M.S.E. and P.J.T. wrote the manuscript with input from all authors.

## Data availability

All data generated or analyzed during this study are included in this article and its supplementary information files. Source data for animal survival cohorts in Figs. 1b, 2c, d, and 4c, i, j and are provided in Supplementary Figs. 1 and 3. Raw annotated western blot images for Extended Data Fig. 1d, f and Extended Data Fig. 7b, c, e are provided as Supplementary Figs. 2 and 4. Animals and iPSC lines are available from P.J.T. upon request.

## Materials and Methods

### Mice

All procedures were in accordance with the National Institutes of Health Guidelines for the Care and Use of Laboratory Animals and were approved by the Case Western Reserve University Institutional Animal Care and Use Committee (IACUC).

Wildtype (B6CBACa-Aw-J/A) and *jimpy* (B6CBACa-Aw-J/A-Plp1jp EdaTa/J) mice used in this study were purchased from Jackson Laboratory (Bar Harbor, ME). *Jimpy* males exhibit severe neurological phenotypes and die around 3 weeks of age due to mutations in *Plp1* on the X-chromosome. The colony was maintained by breeding heterozygous females, which lack a phenotype, to wild-type males to generate affected *jimpy* males. Mice were housed under a temperature-controlled environment, 12-h light-dark cycle with ad libitum access to water and rodent chow. All mice were genotyped at ~postnatal day 7 using genomic DNA isolated from tail tips or toes at two loci: 1) the *jimpy* mutation (NM_011123.4:c.623 -2A>G) in *Plp1* intron 4, which causes skipping of exon 5 and a truncated PLP protein and 2) the complex indel in *Plp1* exon 3 from dual cutting of CRISPR/spCas9 sgRNAs in “CR-*impy*” mice (c.[242_318del; 328_330del]). This “knockdown” causes a frameshift in *Plp1*, a premature stop codon in exon 4, and is predicted to cause nonsense mediated decay of the transcript and loss of protein. Genotyping was performed by standard Sanger sequencing or a custom real time PCR assays (Probe identifiers: Plp1-2 Mut [for *jimpy* mutation in intron 4] and Plp1-5 WT [for *CR-*impy** complex deletion in exon 3], Transnetyx, Cordova, TN).

Primers for Sanger sequencing included:

*jimpy* Forward: AACGCAAAGCAGCACATTTCA

*jimpy* Reverse: AGTGCAGCTCTGGGGTTAAT

CR-*impy* Forward: TCTGTCTGTCCATGCAGGATT

CR*-impy* Reverse: GACACACCCGCTCCAAAGAA

### *Plp1*-targeting sgRNA design

Mouse *Plp1* sequence was entered into the spCas9 CRISPR sgRNA design tool at crispr.mit.edu ^46^ and analyzed against the mm10 target genome. *Plp1*-targeting sgRNAs were sorted based on their on-target efficiency while minimizing off-target mutations. On-target nuclease activity was confirmed for each sgRNA using the Guide-it sgRNA Screening Kit (631440, Clontech) according to the manufacturer’s instructions. The following sgRNAs were tested, and sgRNAs 3 and 7 were selected for combined use in zygote studies based on localization within the gene, proximity to each other, and ability to target *Plp1*, including its splice isoform *Dm20*:

sgRNA1: CCCCTGTTACCGTTGCGCTC
sgRNA2: TGGCCACCAGGGAAGCAAAG
sgRNA3: AAGACCACCATCTGCGGCAA
sgRNA4: GGCCTGAGCGCAACGGTAAC
sgRNA5: GCCTGAGCGCAACGGTAACA
sgRNA6: TCTACACCACCGGCGCAGTC
sgRNA7: CCAGCAGGAGGGCCCCATAA
sgRNA8: GAAGGCAATAGACTGACAGG

### Knockdown of *Plp1* in *jimpy* zygotes using CRISPR-Cas9

Carrier female oocyte donors were administered 5 IU pregnant mare’s serum gonadotropin by intraperitoneal injection (G4877, Sigma-Aldrich), followed by 2.5 IU human chorionic gonadotropin (GC10, Sigma-Aldrich) 48 hours later. These superovulated females were mated to wild-type males. Zygotes were harvested in FHM medium (MR-025 Sigma-Aldrich) with 0.1% hyaluronidase (H3501, Sigma-Aldrich) and the surrounding cumulus cells were separated. The zona pellucida of each zygote was partially dissected using 0.3M sucrose (S7903, Sigma-Aldrich) in FHM as previously described ^47^

Zygotes were placed in 2x KSOM medium (MR-106, Sigma-Aldrich) with an equal volume of solution containing 100ng/uL sgRNA3, 100ng/uL sgRNA7 (AR01, PNAbio), and 200ng/uL spCas9 mRNA (CR01, PNAbio). Electroporation was performed in a chamber with a 1mm gap between two electrodes using an ECM 830 Square Wave Electroporation System (BTX). Electroporation parameters were set as follows: 32V, 3ms pulse duration, 5 repeats, and 100ms inter-pulse interval. Electroporated zygotes were moved to KSOM medium and then transferred into the oviducts of pseudopregnant females (CD1). Electroporation settings were optimized to achieve maximal cutting efficiency in a separate strain but resulted in a higher rate of embryo loss in our B6CBACa/J strain. Zygotes were electroporated in batches of 54, 56, and 61, which resulted in 4, 3, and 0 pups born. The 7 surviving mice were genotyped after birth and monitored daily for onset of typical *jimpy* phenotypes including tremors, seizures, and early death by postnatal day 21. A founder *jimpy* male with complex deletion containing 80-bp of total deleted sequence in exon 3 of *Plp1*, denoted “CR-*impy*” for <u>CR</u>ISPR frameshift-mediated knockdown of *Plp1* in *jimpy*, showed no overt phenotype and was backcrossed for two generations to the wild-type parental strain to reduce potential off-target Cas9 cutting effects (Extended Data Fig. 1b). A colony of mice was bred to evaluate cellular, molecular, and functional phenotypes of contemporaneous isogenic wild-type, *jimpy*, and CR-*impy* male mice. Mice were monitored daily to determine lifespan with statistical significance among groups determined using the log-rank test. Additionally, animals surviving beyond 3 weeks were analyzed using behavioral (rotarod and open field testing for motor performance), histology (immunostaining of the CNS for myelin proteins and electron microscopy for myelin ultrastructure), and electrophysiology (conduction velocity of the optic nerve). Details and metadata for all mice in this study including censoring of animals in the survival analysis are found in Supplementary Fig. 1.

### CRISPR on- and off-target assessment

CRISPR on- and off-target cutting efficiencies were assessed by high throughput sequencing. PCR primers were designed to encompass each guide on-target site, as well as each top predicted off-target site from the spCas9 CRISPR sgRNA design tool at crispr.mit.edu ^46^. Primer sequences were generated using NCBI Primer-BLAST:

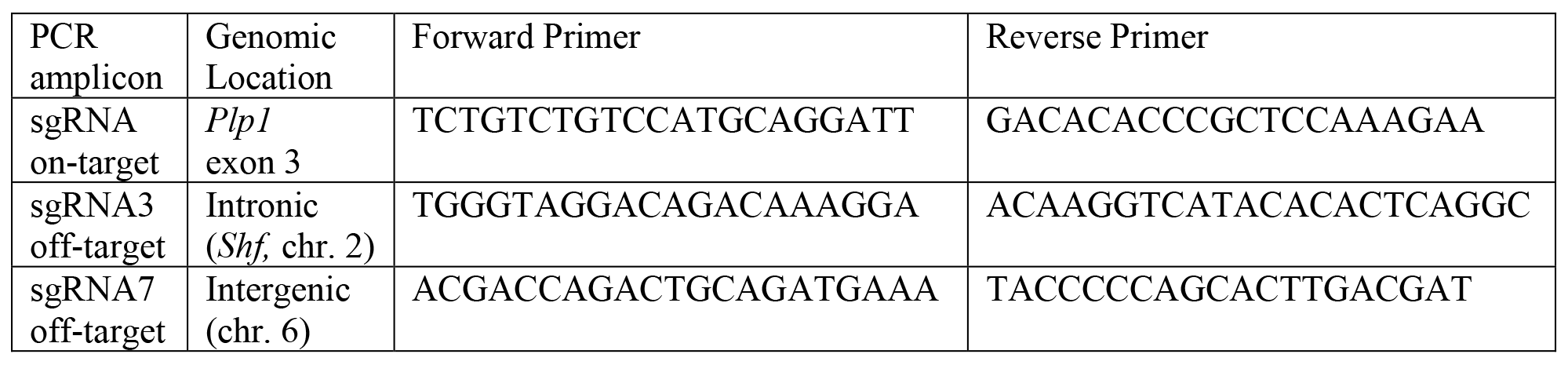

Tails were added to each primer sequence:

Forward: TCCCTACACGACGCTCTTCCGATCT

Reverse: AGTTCAGACGTGTGCTCTTCCGATCT

PCR amplification was performed using the KAPA HiFi HotStart ReadyMix (07958935001, Roche) to minimize PCR-based error. Libraries were prepared by adding unique indices by PCR using KAPA HiFi HotStart ReadyMix. All libraries were pooled evenly and quantified using NEBNext^®^ Library Quant Kit for Illumina^®^ (E7630, New England Biolabs) then denatured and diluted per Illumina’s MiSeq instructions. 250bp paired-end sequencing was performed using an Illumina MiSeq at the Case Western Reserve University School of Medicine Genomics Core Facility. Reads were compared against the consensus sequence and CRISPR-induced indel percentages were determined using the OutKnocker tool at outknocker.org ^48^.

### Video recording of mouse phenotypes

All recording was performed using video recording function on an iPhone (Apple). Videos were color corrected, stabilized, and trimmed to a discrete range using iMovie (Apple). Videos were collated and converted to MP4 format using Adobe After Effects.

### Immunohistochemistry

Mice were anesthetized with isoflurane and sacrificed by transcardial perfusion with PBS followed by 4% paraformaldehyde (PFA; 15710, Electron Microscopy Sciences). Tissue was harvested and placed in 4% PFA overnight at 4°C. Samples were rinsed with PBS, equilibrated in 30% sucrose, and frozen in Tissue-Tek^®^ Optimum Cutting Temperature compound (O.C.T.; 25608-930, VWR). Samples were cryosectioned at a 20μm thickness. Sections were washed in phosphate-buffered saline (PBS) and incubated overnight in antibody solution containing 2.5% normal donkey serum (NDS; 017-000-121, Jackson Labs) and 0.25% Triton X-100 (Sigma: T8787). For MBP immunohistochemistry, sections were post fixed in methanol at – 20°C for 20 minutes followed by overnight incubation in a PBS based primary antibody solution containing 0.1% Saponin and 2.5% normal donkey serum. Sections were stained using the following antibodies at the indicated concentrations or dilutions: mouse anti-MBP (2μg/mL, 808401, Biolegend), rabbit anti-MyRF antibody (1:500; kindly provided by Dr. Michael Wegner), rat anti-PLP (1:500; Lerner Research Institute Hybridoma Core, Cleveland, OH), goat anti-Sox10 (0.4μg/mL, AF2864, R&D Systems), rabbit anti-GFAP (1:1000; Z033429-2, Dako), goat anti-IBA1 (0.1mg/mL, ab5076, Abcam), anti-phospho-neurofilament (2ug/ml, 801601, BioLegend), and rabbit anti-ASO (1:2500; Ionis Pharmaceuticals). Secondary immunostaining was performed with Alexa Fluor^®^ antibodies (ThermoFisher) used at 1ug/ml. Nuclei were identified using 100ng/ml DAPI (D8417, Sigma). Stained sections were imaged using the Operetta^®^ High Content Imaging and Analysis system (PerkinElmer) and Harmony^®^ software (PerkinElmer) for whole section images and a Leica Sp8 confocal microscope or a Leica DMi8 inverted microscope with Leica Application Suite X software for all other immunohistochemical imaging. To quantify MyRF staining, MyRF+ cells were counted along the length of the whole corpus callosum from medial sagittal sections from three animals per genotype. A one-way ANOVA with correction for multiple comparisons or a Welch’s t-test was performed to determine statistical significance across genotypes.

### qRT-PCR

Mice were euthanized using isoflurane overdose. Different brain regions (cerebral cortex, cerebellum, and brainstem) were harvested. Each brain region was split in two and half was used for RNA quantification using qRT-PCR, the other for western blot analysis (see below). TRI Reagent (R2050-1-200, Zymo Research) was separately added to tissue and samples were homogenized using Kontes Pellet Pestle Grinders (KT749520-0000, VWR). RNA was extracted using the RNeasy Mini Kit (74104, Qiagen) according to the manufacturer’s instructions. Reverse transcription was performed using the iScript cDNA Synthesis Kit (1708891, Biorad) with 1μg of RNA per reaction. Real-Time PCR was then performed on an Applied Biosystems 7300 Real-time PCR system with 10ng cDNA per sample in quadruplicate using Taqman gene expression master mix (4369016, ThermoFisher) and the following pre-designed Taqman gene expression assays (4351370, ThermoFisher): *Plp1* (Mm01297210_m1), *Mbp* (Mm01266402_m1) and *Actb* (Mm00607939_s1) (endogenous control). Expression values were normalized to *Actb* and to wild-type samples (for CRISPR cohort) or wild-type untreated samples (for ASO-treated wild-type cohort). Graphpad Prism software was used to perform a one-way ANOVA with Tukey correction or a one-way ANOVA with Dunnett’s correction for multiple comparisons to determine statistical significance across genotypes or ASO treatments, respectively.

### Protein quantification and western blot

Tissues were obtained as described above. Protein lysis buffer consisting of RIPA buffer (R0278, Sigma), cOmplete™ Mini EDTA-free Protease Inhibitor Cocktail (11836170001, Sigma), Phosphatase Inhibitor Cocktail 3 (P0044, Sigma), Phosphatase Inhibitor Cocktail 2 (P5726, Sigma), and BGP-15 (B4813, Sigma) was added to each sample. Tissue was homogenized using Dounce Tissue Grinders (D8938, Sigma). Lysate was separated by centrifugation at 17000g for 15 minutes at 4°C. A BCA standard curve was generated using the Pierce BCA Protein Assay Kit (23225, Thermo Scientific) and used to samples to an equivalent protein concentration. Equal amounts of sample were run on a NuPAGE 4-12% Bis-Tris Protein gel (NP0335BOX or NP0329BOX, Thermo Fisher), then electrophoretically transferred to a PVDF membrane (LC2002, Invitrogen or 926-31097, Li-Cor). The membrane was blocked with 5% milk in TBS-T for an hour, then hybridized with mouse anti-MBP antibody (1μg/mL; 808401, Biolegend) or rat anti-PLP antibody (1:1000; Lerner Research Institute Hybridoma Core, Cleveland, OH) overnight at 4°C. Blots were then washed in TBS-T and incubated in goat antimouse HRP (1:2500, 7076, Cell Signaling), goat anti-rat HRP (1:2500, 7077, Cell Signaling), or IRDye secondaries (1:20000, 925, Li-Cor). Each sample was normalized to B-actin using HRP-conjugated mouse anti-B-actin (1:10000, A3854-200UL, Sigma-Aldrich). All secondary antibodies were incubated for one hour at room temperature. Blots were analyzed with the Odyssey^®^ Fc imaging system (Li-Cor). Graphpad Prism software was used to perform a one-way ANOVA with Tukey correction or a one-way ANOVA with Dunnett’s correction for multiple comparisons to determine statistical significance across genotypes or ASO treatments, respectively. Raw annotated images of full western blots are provided in Supplementary Figs. 2 and 4.

### Electron microscopy and toluidine blue staining

Mice were anesthetized with isoflurane and rapidly euthanized. Tissue was collected after transcardial perfusion with PBS followed by 4% paraformaldehyde and 2% glutaraldehyde (16216, Electron Microscopy Sciences) in 0.1M sodium cacodylate buffer, pH 7.4 (11652, Electron Microscopy Sciences), except for 6 month optic nerve samples which were placed directly into fixative without perfusion. Samples were post-fixed with 1% osmium tetroxide (19150, Electron Microscopy Sciences) and stained with 0.25% uranyl acetate (22400, Electron Microscopy Sciences), en bloc. Samples were dehydrated using increasing concentrations of ethanol, passed through propylene oxide, and embedded in Eponate 12™ epoxy resin (18012, Ted Pella). Silver-colored sections were prepared (Leica EM UC6), placed on 300 mesh nickel grids (T300-Ni, Electron Microscopy Sciences), stained with 2% uranyl acetate in 50 % methanol, and stained with lead citrate (17800, Electron Microscopy Sciences). Sections were imaged using a FEI Tecnai Spirit electron microscope at 80 kV. Myelinated axons were manually counted from the sections made on the middle part of the nerve lengthwise and at least three areas across the optic nerve diameter using Adobe Photoshop (Adobe Systems). Graphpad Prism software was used to perform a one-way ANOVA with Tukey correction to determine significance across genotypes. Toluidine Blue (22050, Electron Microscopy Sciences) stained 1μm sections were prepared from same epoxy resin embedded samples above and visualized with a light microscope (Zeiss Axioskop2) using plan-NEOFLUAR 100X 1.30 oil objective lens and images were captured using a Scion 1394 color camera with ImageJ software.

### Optic nerve electrophysiology

Mice were deeply anesthetized with isoflurane and euthanized. Each eye with its attached optic nerve was dissected and placed in Ringer’s solution consisting of 129mM NaCl (BP358-212, Fisher Scientific), 3mM KCl (BP366-500, Fisher Scientific), 1.2mM NaH_2_PO_4_ (1-3818, J. T. Baker Chemical), 2.4mM CaCl_2_ (C79-500, Fisher Scientific), 1.3mM MgSO_4_ (M2643, Sigma), 20mM NaHCO_3_ (S233-500, Fisher Scientific), 3mM HEPES (H3375, Sigma), 10mM glucose (G5767, Sigma), oxygenated using a 95%O_2_/5% CO_2_ gas mixture. Each nerve was carefully cleaned, transected behind the eye, at the optic chiasm, and allowed to recover for one hour in oxygenated Ringer’s solution at room temperature (22-24°C). Each end of the nerve was set in suction electrodes, pulled from polyethylene tubing (PE-190, BD Biosciences). Monophasic electrical stimuli were applied to the proximal end of the nerve and recordings were captured at the distal end. The recovery of the response was monitored every 20 min for one hour, and only fully recovered samples were subjected to additional stimuli. Stimuli were generated with a S48 stimulator (Grass Technologies) and isolated from ground with PSIU6B unit (Grass Technologies). Supra-threshold stimulus was determined using 30μs stimulus duration. The response was amplified 100X with P15D preamplifier (Grass Technologies), monitored with oscilloscope (V1585, Hitachi), digitized with Digidata1550A (Axon Instruments) and recorded using 50kHz sampling rate with AxoScope software (Axon Instruments). The distance between the electrodes was measured and used to calculate the conduction velocity of the compound action potential (CAP) peaks at their latency. Recorded signals were analyzed using AxoScope software.

### Open Field Testing

Locomotion was assessed by open field testing. Animals were placed in the center of a 20-inch by 20-inch square box and all movements were captured for a total of five minutes using ANY-maze software version 5.0 (Stoelting Co). Total distance traveled was reported for each animal. Graphpad Prism software was used to perform a one-way ANOVA with Tukey correction or a one-way ANOVA with Dunnett’s correction for multiple comparisons to determine statistical significance across genotypes or ASO treatments, respectively.

### Rotarod Testing

Motor performance was assessed using a Rota Rod Rotomax 5 (Columbus Instruments) with a 3cm diameter rotating rod. Immediately prior to testing animals were trained at a constant speed of 4 rounds per minute (rpm) for a total of two minutes. Testing began at 4 rpm with an acceleration of 0.1 rpm/s. Time to fall was recorded from three independent trials, and the average value for each animal was reported. Between training and each experimental trial animals were allowed to rest for at least five minutes. Graphpad Prism software was used to perform a oneway ANOVA with Tukey correction or a one-way ANOVA with Dunnett’s correction for multiple comparisons to determine statistical significance across genotypes or ASO treatments, respectively.

### Generation of iPSCs

Tail tips (2 mm piece from 8 day old CR-*impy* mice) were bisected, placed on Nunclon-Δ 12-well plates (150628, ThermoFisher), and covered with a circular glass coverslip (12-545-102; Fisher Scientific) to maintain tissue contact with the plate and enable fibroblast outgrowth. Tail-tip fibroblasts were cultured in ‘fibroblast medium’ consisting of DMEM (11960069, ThermoFisher) with 10% fetal bovine serum (FBS; 16000044, ThermoFisher), 1x nonessential amino acids (11140050, ThermoFisher), 1x Glutamax (35050061, ThermoFisher), and 0. 1 mM 2-mercaptoethanol (M3148, Sigma Aldrich) supplemented with 100U/mL penicillin-streptomycin (15070-063, ThermoFisher). Medium was changed every day for the first 3 days and then every other day.

Fibroblasts were seeded at approximately 1.4×10^4^ cells/cm^2^ on Nunclon-Δ dishes in fibroblast medium, and allowed to equilibrate overnight. The following day medium was removed and replaced with an equal volume of pHAGE2-TetOminiCMV-STEMCCA-W-loxp lentivirus encoding a floxed, doxycycline-inducible polycistronic Oct4, Sox2, Klf4, and c-Myc construct and pLVX-Tet-On-Puro (632162, Clontech) lentivirus supplemented with 8μg/mL polybrene (107689, Sigma). Lentivirus was prepared using the Lenti-X Packaging Single Shots (631275, Clontech) according to manufacturer’s instructions. Three hours later lentivirus medium was removed and replaced with fibroblast medium supplemented with 2 μg/ml doxycycline (631311, Clontech). The following day media was removed and replaced with an equal volume of pHAGE2-TetOminiCMV-STEMCCA-Wloxp and pLVX-Tet-On-Puro lentivirus supplemented with 8μg/mL polybrene. Three hours later lentivirus media was diluted 1:2 with fibroblast medium. Medium was changed each day with fibroblast medium supplemented with 2 μg/ml doxycycline and 10^3^ units/ml LIF. After 3 days fibroblasts were lifted using Accutase and seeded on Nunclon-Δ plates, atop a feeder layer of irradiated mouse embryonic fibroblasts (iMEFs; produced in-house) previously plated at 1.7×10^4^ cells/cm^2^ on 0.1% gelatin (1890, Sigma) coated Nunclon-Δ plates in “pluripotency medium” consisting of Knockout DMEM (10829-018, ThermoFisher), 5% FBS, 15% knockout replacement serum (10828028, ThermoFisher), 1x Glutamax, 1x nonessential amino acids, 0.1 mM 2-mercaptoethanol, and 10^3^ units/ml LIF (LIF; ESG1107, EMD Millipore) supplemented with 2 μg/ml doxycycline. Medium was changed every day until iPSC colonies began to emerge. Individual colonies were picked and dissociated in accutase and were individually plated in single wells of Nunclon-Δ 12-well plates, atop an iMEF feeder layer in pluripotency medium supplemented with 2 μg/ml doxycycline. Clones were further expanded, with daily medium changes. iPSC colonies were stained for pluripotency markers and karyotyped at the seventh passage after derivation (Cell Line Genetics; Madison, WI). CR-*impy* iPSCs were derived and characterized for this study (line identifier jpCR100.1). Isogenic comparator *jimpy* (line identifier i.jp-1.6) and wild-type (line identifier i.wt-1.0) iPSC lines were described and characterized separately ^22^. Genotypes of iPSCs were re-verified prior to use.

### Generation of iPSC-derived OPCs

iPSCs were differentiated to OPCs as previously described ^25,26^. In brief, iPSC were isolated from their iMEF feeder layer using 1.5mg/mL collagenase type IV (17104019, ThermoFisher) and dissociated with either 0.25% Typsin-EDTA or Accutase and seeded at 7.8×10^4^ cells/cm2 on Costar Ultra-Low attachment 6-well plates (3471, Corning). Cultures were then directed through a stepwise differentiation process to generate pure populations of OPCs. OPCs were maintained in “OPC medium” consisting of DMEM/F12 (11320082, ThermoFisher), 1x N2 supplement (AR009, R&D Systems), 1x B-27 without vitamin A supplement (12587-010, ThermoFisher), and 1x Glutamax (collectively “N2B27 medium”), supplemented with 20 ng/mL fibroblast growth factor 2 (FGF2; 233-FB, R&D Systems) and 20 ng/mL platelet-derived growth factor-AA (PDGF-AA; 221-AA, R&D Systems). Medium was changed every other day. For characterization of purity, iPSC-derived OPCs from all genotypes were fixed with 4% PFA and immunostained for canonical OPC transcription factors, Olig2 and Sox10.

### *In vitro* assessment of oligodendrocyte differentiation and survival from OPCs

OPCs from each genotype were plated in parallel onto Nunclon-Δ 96-well plates (150628, ThermoFisher) that were first coated with 100 μg/mL poly(L-ornithine) (P3655, Sigma), followed by 10 μg/ml laminin solution (L2020, Sigma). For the oligodendrocyte differentiation assay, 25,000 cells were seeded per well in media that consisted of DMEM/F12 (11320082, ThermoFisher), 1x N2 supplement (AR009, R&D Systems), 1x B-27 without vitamin A supplement (12587-010, ThermoFisher), and 1x Glutamax, supplemented with T3 (40ng/ml), Noggin (100ng/ml), cAMP (10uM), IGF (100ng/ml) and NT3 (10ng/ml). All plates were incubated at 37°C and 5% CO_2_ for 3 days. Cells were immunostained using for myelin protein markers. Eight fields were captured per well and the total number of MBP+ cells were quantified for each cell line. Graphpad Prism software was used to perform a one-way ANOVA with Tukey correction for multiple comparisons to determine statistical significance across genotypes.

### Immunocytochemistry

Cells were fixed with 4% paraformaldehyde (PFA) in phosphate buffered saline (PBS). After fixation, cells were permeabilized with 0.2% Triton X-100 in PBS followed by blocking in 10% donkey serum in PBS. Cells were stained overnight at 4°C with the following primary antibodies diluted in blocking solution: rat anti-MBP (1:100; ab7349, Abcam), rat anti-PLP (1:5000; Lerner Research Institute Hybridoma Core, Cleveland, OH), goat anti-Sox10 (2μg/mL; AF2864, R&D Systems), rabbit anti-Olig2 (1:1000; 13999-1-AP, ProteinTech), rabbit anti-Nanog (0.4μg/mL; AB21624, Abcam), mouse anti-Oct4 (0.4μg/mL; SC-5279, Santa Cruz). For secondary immunostaining, Alexa Fluor^®^ antibodies (ThermoFisher) were used at 1ug/ml, and DAPI (100ng/mL, D8417, Sigma) was used at to identify nuclei.

### ASO design and characterization

Second generation ASOs were designed to target mouse *Plp1*. ASOs consisted of 20-mer nucleotide sequences with 2’-O-methoxyethyl (MOE) modifications and a mixed backbone of phosphorothioate and phosphodiester internucelotide linkages.

ASOs were screened for efficacy in primary E16 cortical cultures, as previously described ^35^. Briefly, cells were treated with ASOs at 37°C/5% CO_2_ for 3 days, RNA was isolated, and *Plp1* transcript level was quantified with qRT-PCR on Step One instruments (Thermo Fisher). *Plp1* mRNA was normalized to total RNA measured with the Quant-iTTM RiboGreen^®^ RNA reagent. ASOs that efficiently reduced *Plpl* mRNA were selected for *in vivo* screening and tolerability studies.

Lead ASOs were administered to 8 week old C57BL/6J mice (Jackson Labs) via single 500 μg intracerebroventricular (ICV) injection and *Plp1* mRNA levels were measured by RT-qPCR in cortex and spinal cord tissue after 2 weeks. ASOs with greater than 90% *Plp1* mRNA reduction were selected for further characterization. These were administered to mice via single 300 μg ICV bolus injection to test for efficacy and tolerability, as measured by markers of glial cell activation, 8 weeks post-ICV. Levels of *Plp1* mRNA as well as markers of astro- or micro-glial activation, *Gfap, Aif1*, and *CD68*, were assessed by RT-qPCR using the following custom primer/probe sets (Integrated DNA Technologies):

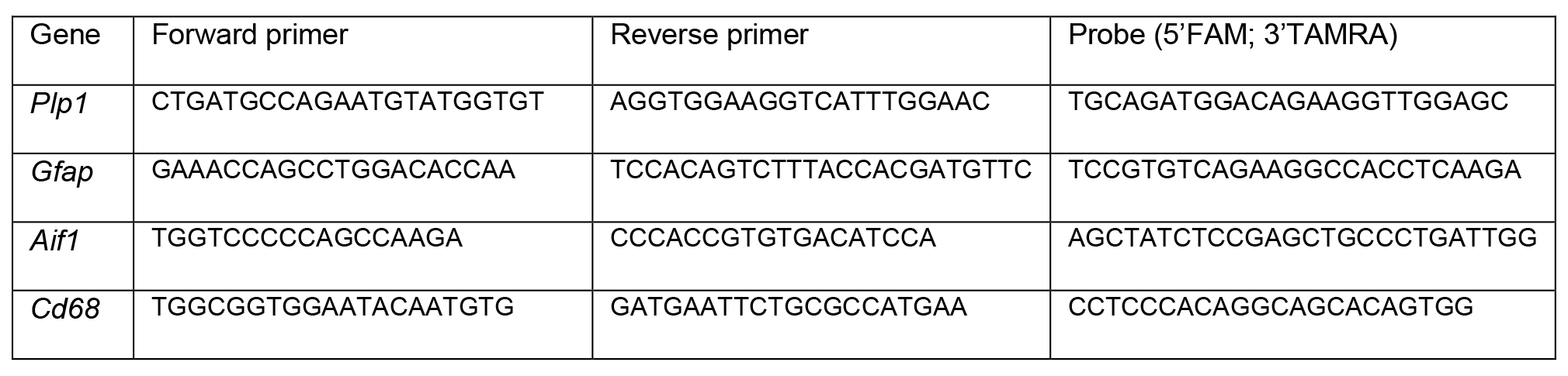

Immunohistochemical staining was used to assess morphology of astrocytes (rabbit polyclonal antibody, DAKO) and microglia (rabbit polyclonal antibody, WAKO) in formalin-fixed, paraffin embedded brain and spinal cord sections. *Plp1* ASO.a (intron 5) and ASO.b (3’ UTR) were selected for use in j*impy* mice, as well as a control ASO with no known murine target.

### Therapeutic application of ASOs to postnatal mice

Male pups from crosses between *jimpy* mutation carrier females and wild-type males were administered 30ug of either *Plp1*-targeting ASOs *Plp1*.a, *Plp1*.b, a control non-targeting ASO, or left untreated. ASOs were administered using a Hamilton 1700 gastight syringe (7653-01, Hamilton Company) by ICV injection to cryoanesthetized mice. The needle was placed between bregma and the eye, 2/5 the distance from bregma, and inserted to a depth of 2mm according to published protocols ^49^. A total volume of 2uL was administered to the left ventricle. Mice were allowed to recover on a heating pad and subsequently reintroduced to the mother.

Mice were genotyped at approximately postnatal day 7 and monitored daily for onset of typical *jimpy* phenotypes including tremors, seizures, and early death by 3 weeks of age. Lifespan was determined for each animal with statistical significance among groups determined using the log-rank test. All mice surviving to a pre-determined endpoint of 8 months of age were sacrificed for histological analysis. Additionally, animals surviving beyond 3 weeks were analyzed using behavioral (rotarod and open field testing for motor performance). Details and metadata for all mice in this study are found in Supplementary Fig. 3.

**Extended Data Fig. 1.**
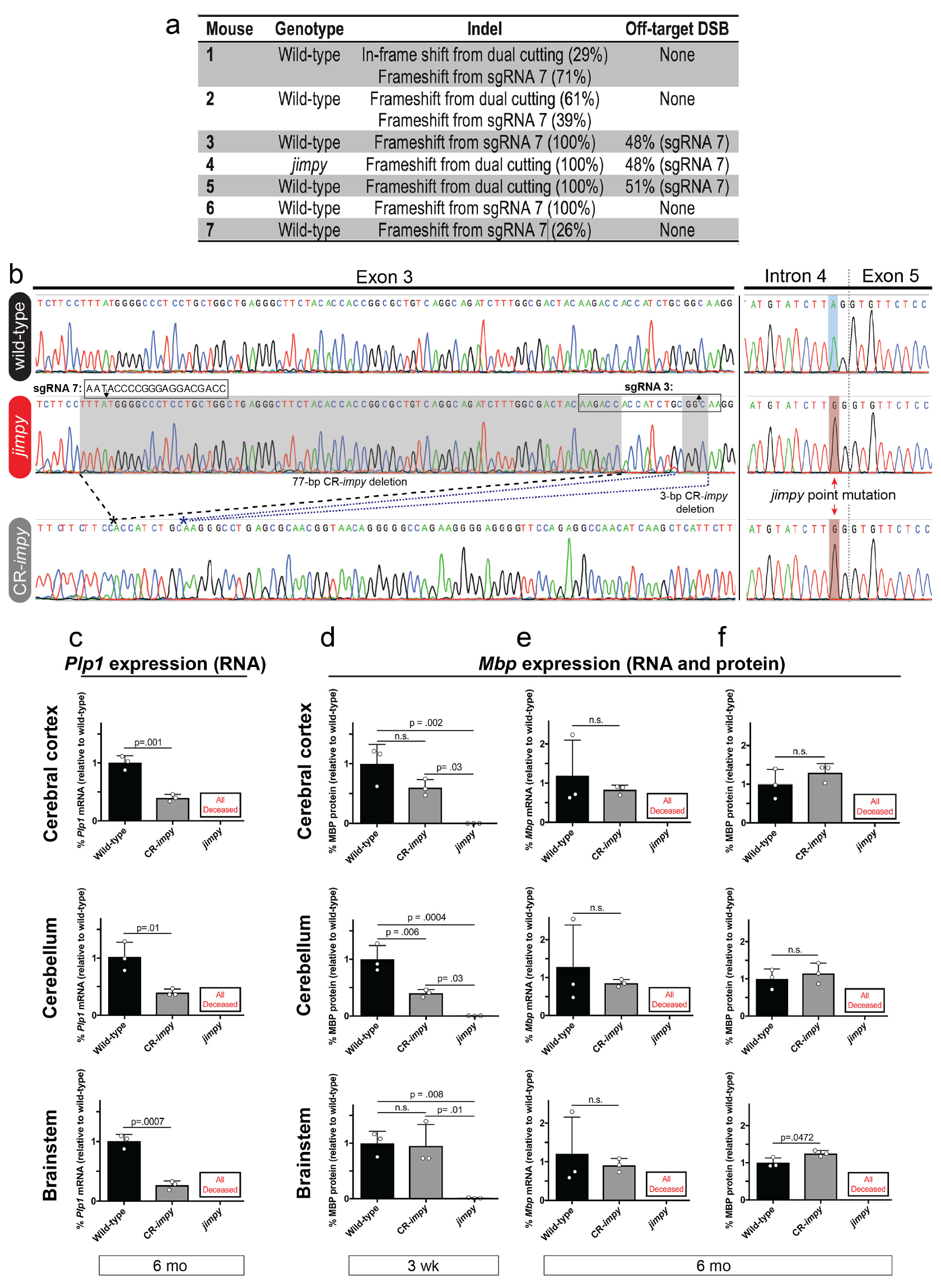
CRISPR-mediated knockdown of *Plp1* in *jimpy* mice increases MBP transcript and protein levels. **a**, Table showing the on- and off-target mutations for sgRNAs 3 and 7 after electroporation into mouse zygotes and measured by high-throughput sequencing of tail tip DNA from founder animals. Mouse number 4, a *jimpy* male with a complex, frameshift deletion including 80-bp of total deleted sequence in *Plp1* exon 3, served as the founder for the CR-*impy* cohort. **b**, Annotated Sanger sequencing traces of wild-type, *jimpy*, and CR-*impy* mice showing the complex, frameshift in *Plp1* exon 3 from dual cutting of CRISPR/spCas9 sgRNAs in CR-*impy* mice as well as the *jimpy* point mutation in intron 4. sgRNA 3 and 7 sequences outlined by black boxes with the predicted double strand break site shown a black arrow. **c**, RT-qPCR data comparing the level of *Plp1* transcript in 6 month old CR-*impy* mice relative to wild-type mice in three different brain regions (cerebral cortex, cerebellum, and brainstem). Primer sites span *Plp1* exons 2-3, upstream of CR-*impy* complex, frameshift deletion and *jimpy* mutation sites. Individual data points represent the mean value of 4 technical replicates for each biological replicate (n=3 separate mice). Error bars show mean ± standard deviation. p-values calculated using a two-way, unpaired t-test. **d**, Western blot data comparing the level MBP in 3 week old CR-*impy* and *jimpy* mice relative to wild-type mice across three different brain regions (cerebral cortex, cerebellum, and brainstem). n=3 biological replicates (separate mice) per genotype (see Supplementary Fig. 2 for full western blot images for all samples). Error bars show mean ± standard deviation. p-values calculated using a one-way ANOVA with Tukey correction for multiple comparisons. **e**, RT-qPCR data comparing the level of *Mbp* transcript in 6 month old CR-*impy* mice relative to wild-type mice in three different brain regions (cerebral cortex, cerebellum, and brainstem). Individual data points represent the mean value of 4 technical replicates for each biological replicate (n=3 separate mice). Error bars show mean ± standard deviation. p-values calculated using a two-way, unpaired t-test. **f**, Western blot data comparing the level MBP in 6 month old CR-*impy* mice relative to wild-type mice in three different brain regions (cerebral cortex, cerebellum, and brainstem). n=3 biological replicates (separate mice) per genotype (see Supplementary Fig. 2 for full western blot images for all samples). Error bars show mean ± standard deviation. p-values calculated using a two-way, unpaired t-test.

**Extended Data Fig. 2.**
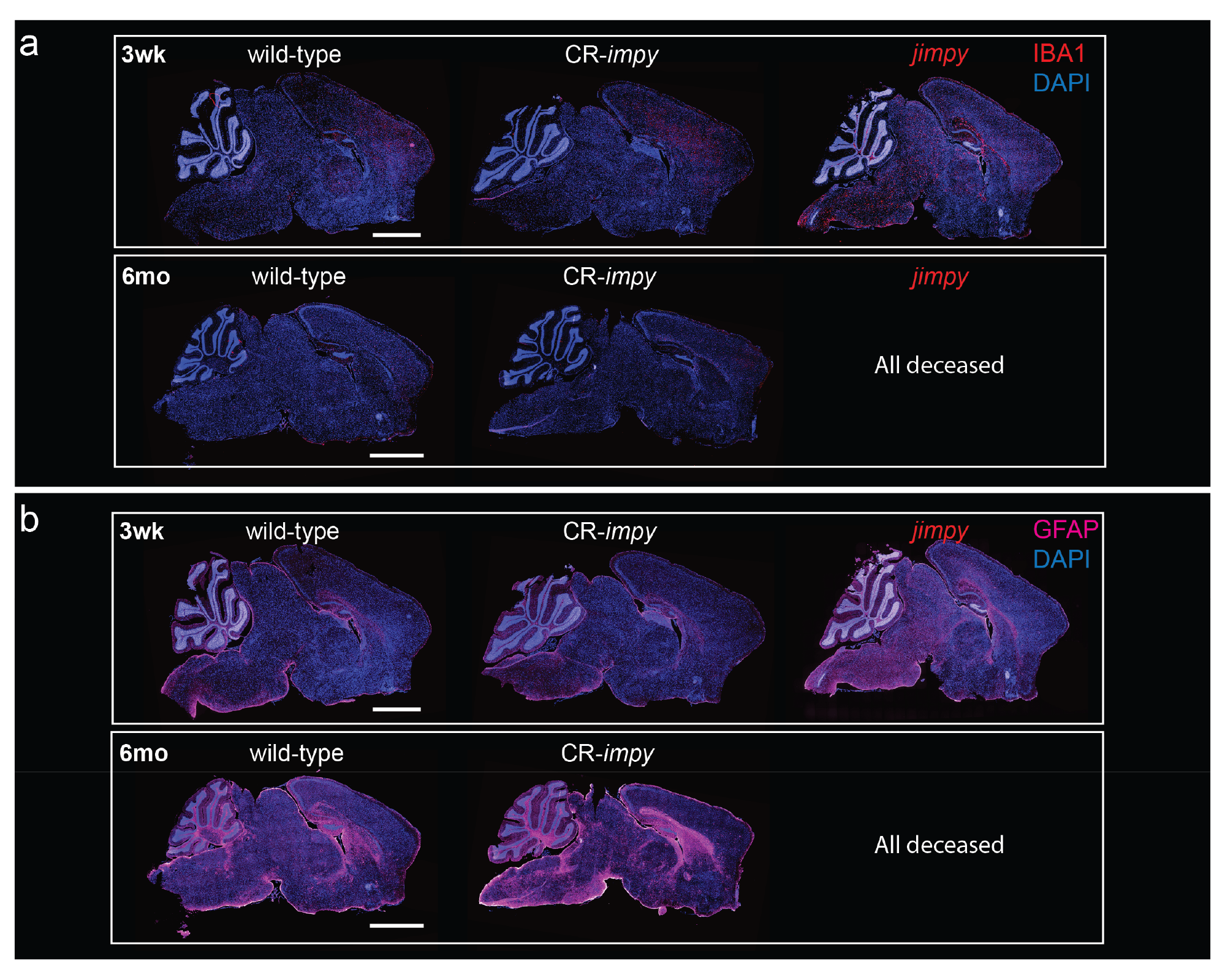
CRISPR-mediated knockdown of *Plp1* in *jimpy* mice reduces activated microglia and astrocyte markers. **a**, Representative immunohistochemical images of whole-brain sagittal sections showing IBA1+ activated microglia (red), and total DAPI+ cells (blue) in wild-type, CR-*impy*, and *jimpy* mice at 3 weeks and 6 months of age as indicated. Scale bar, 2mm. **b**, Representative immunohistochemical images of whole-brain sagittal sections showing GFAP+ astrocytes (magenta), and total DAPI+ cells (blue) in wild-type, CR-*impy*, and *jimpy* mice at 3 weeks and 6 months of age as indicated. Scale bar, 2mm.

**Extended Data Fig. 3.**
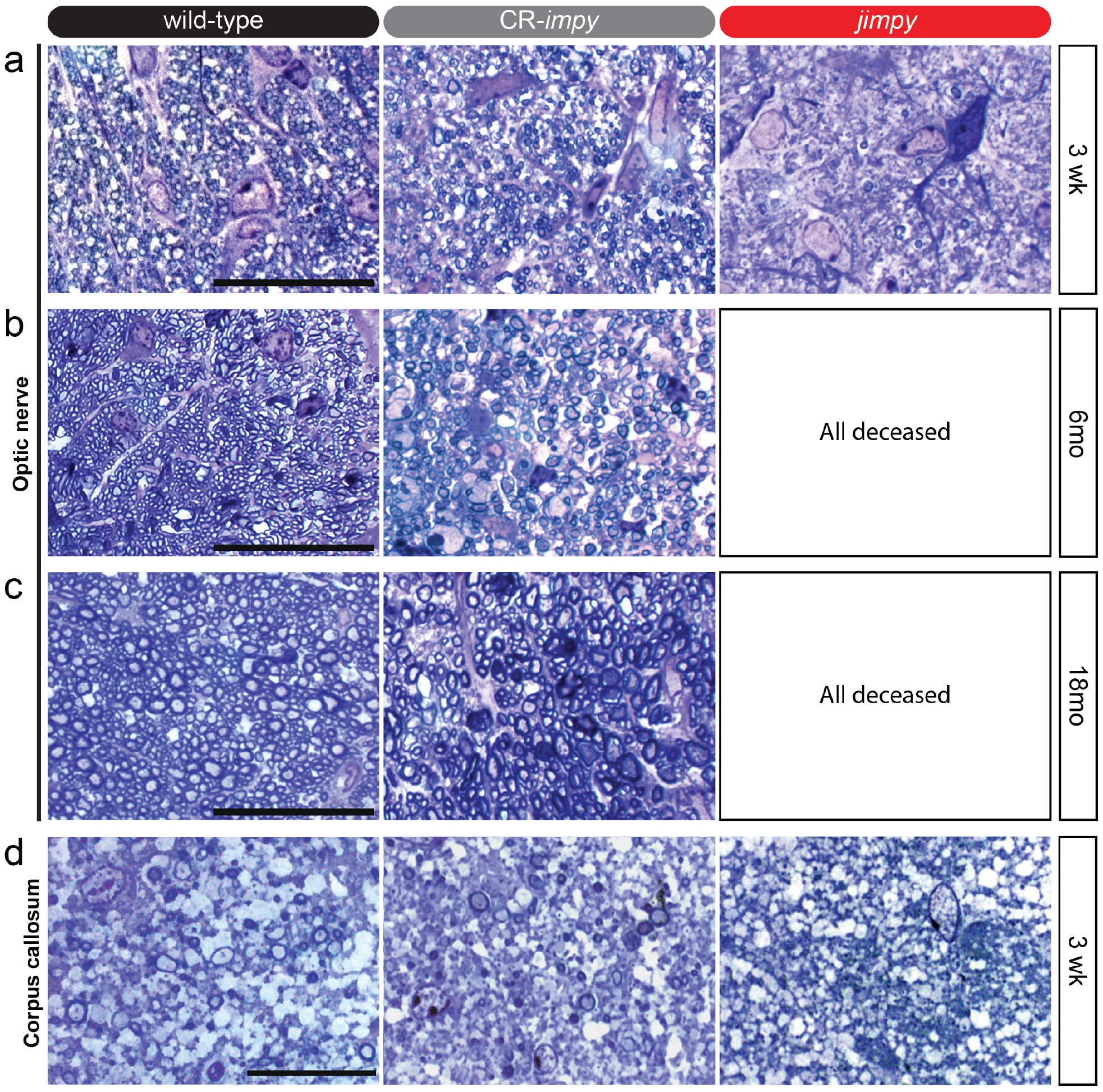
CRISPR-mediated knockdown of *Plp1* in *jimpy* mice increases myelination. **a**, Representative images of toluidine blue-stained optic nerve sections from 3 week old wild-type, *CR-*impy**, and *jimpy* mice. Scale bar, 20μm. **b-c**, Representative images of toluidine blue-stained optic nerve sections from 6 month (b) and 18 month (c) old wild-type, *CR-*impy**, and *jimpy* mice. Scale bar, 20μm. **d**, Representative images of toluidine blue-stained corpus callosum sections from 3 week old wild-type, *CR-*impy**, and *jimpy* mice. Scale bar, 20μm.

**Extended Data Fig. 4.**
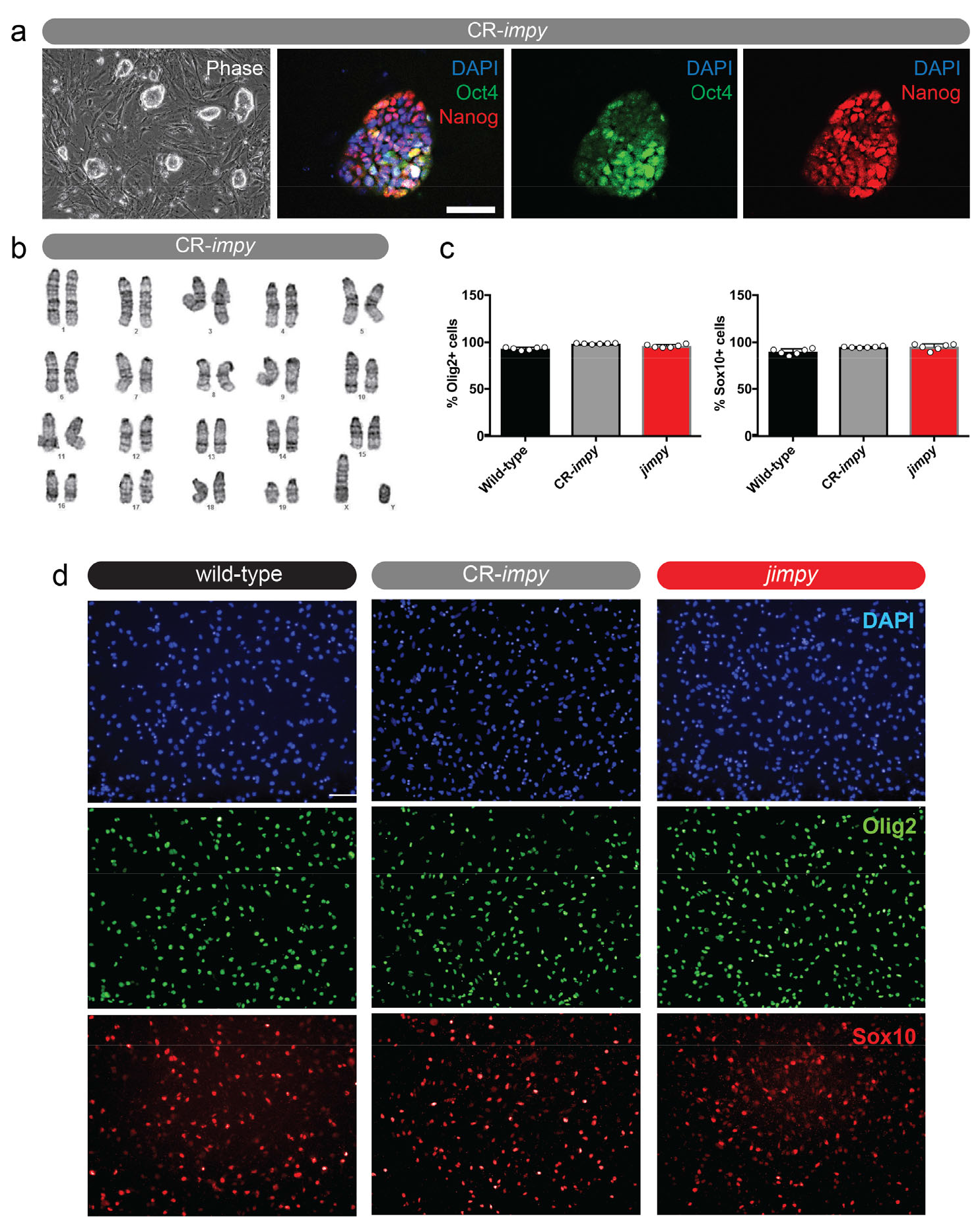
Characterization of mouse iPSC lines and derivation of OPCs. **a**, Representative phase and immunocytochemistry images of Oct4+ (green) and Nanog+ (red) iPSCs reprogrammed from CR-*impy*, tail-tip fibroblasts. Scale bar, 50um. **b**, Normal karyotype of CR-*impy* iPSC line used to generate OPCs. **c**, Percentage of Sox10+ and Olig2+ cells in OPCs cultures from wild-type, CR-*impy*, and *jimpy* iPSCs. Error bars show mean ± standard deviation. n=6 technical replicates (single cell line per genotype with 6 separate wells scored). **d**, Representative immunocytochemistry images showing relative purity of Olig2+ and Sox10+ OPCs derived from wild-type, CR-*impy*, and *jimpy* iPSCs. Scale bar, 100um.

**Extended Data Fig. 5.**
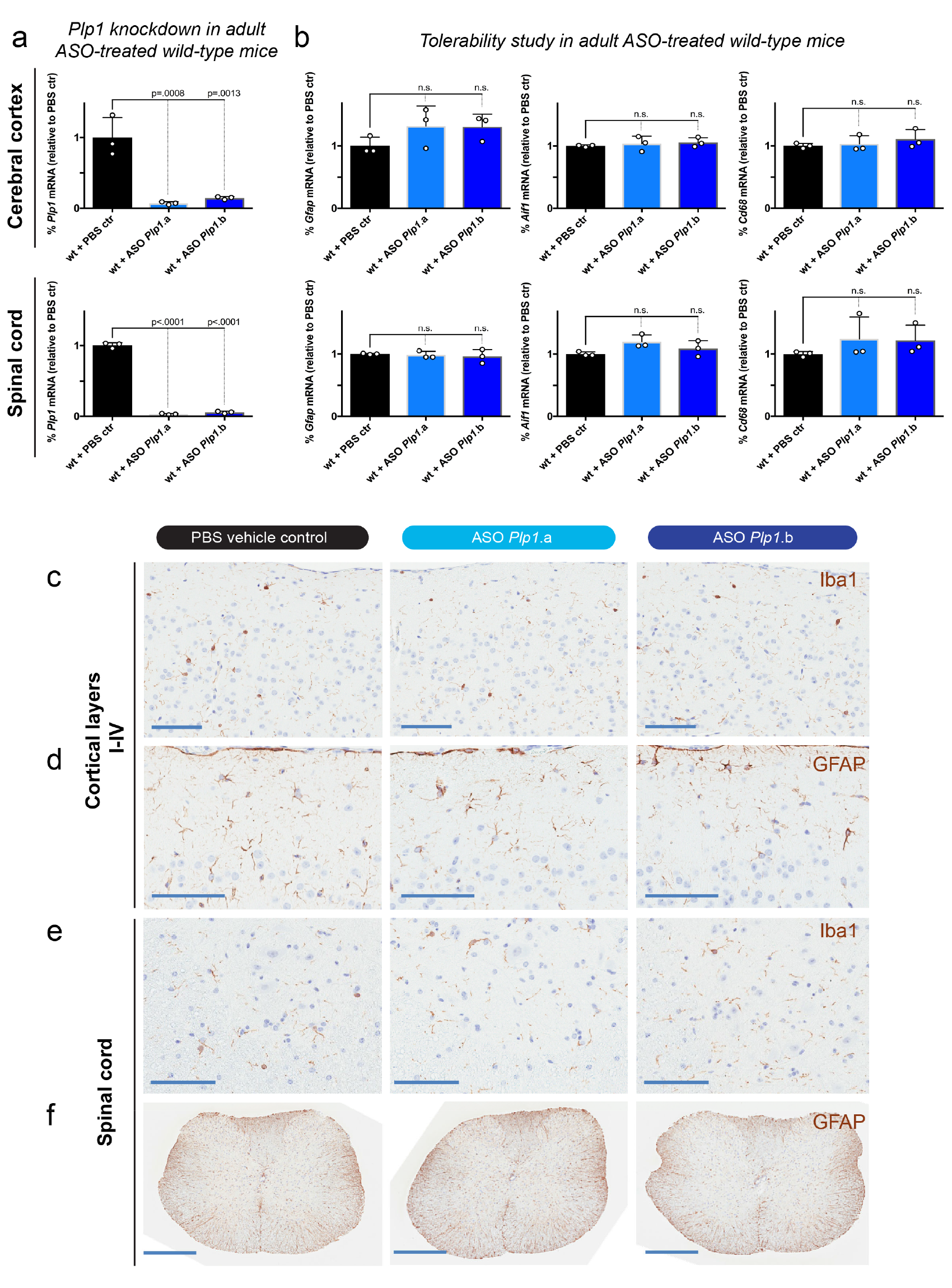
*Plp1*-targeted ASOs show robust and sustained *Plp1* suppression and do not alter markers of glial activation/recruitment in wild-type adult mouse CNS. **a**, RT-qPCR data comparing the level of *Plp1* transcript suppression in 16 week old cerebral cortex and spinal cord of wild-type mice treated with the indicated ASOs (300ug dose) or controls at week 8. Individual data points represent the mean value of 3 technical replicates for each biological replicate (n=3 separate mice). Error bars show mean ± standard deviation. p-values calculated using one-way ANOVA with Dunnett’s correction for multiple comparisons. **b**, RT-qPCR data assessing ASO tolerability by expression levels of *Gfap* (astrocyte marker), *Aifl* (microglia marker), *and Cd68* (monocyte/macrophage marker) transcripts in 16 week old cerebral cortex and spinal cord of wild-type mice treated with the indicated ASOs (300ug dose) or controls at week 8. Individual data points represent the mean value of 3 technical replicates for each biological replicate (n=3 separate mice). Error bars show mean ± standard deviation. p-values calculated using one-way ANOVA with Dunnett’s correction for multiple comparisons. **c-f**, Ibal or GFAP immunohistochemistry (brown) with hematoxylin counterstain (purple) showing no appreciable increase among groups in staining intensity, cellularity, or shortened, thick processes that would be consistent with glial activation. (c) cortical layers I-IV Ibal staining, scale bar = 100 μm (d) cortical layers I-III GFAP staining, scale bar = 100 μm. (e) spinal cord dorsal horn grey/white matter intersection Ibal staining, scale bar = 100 μm. (f) Spinal cord GFAP staining, scale bar = 500 μm.

**Extended Data Fig. 6.**
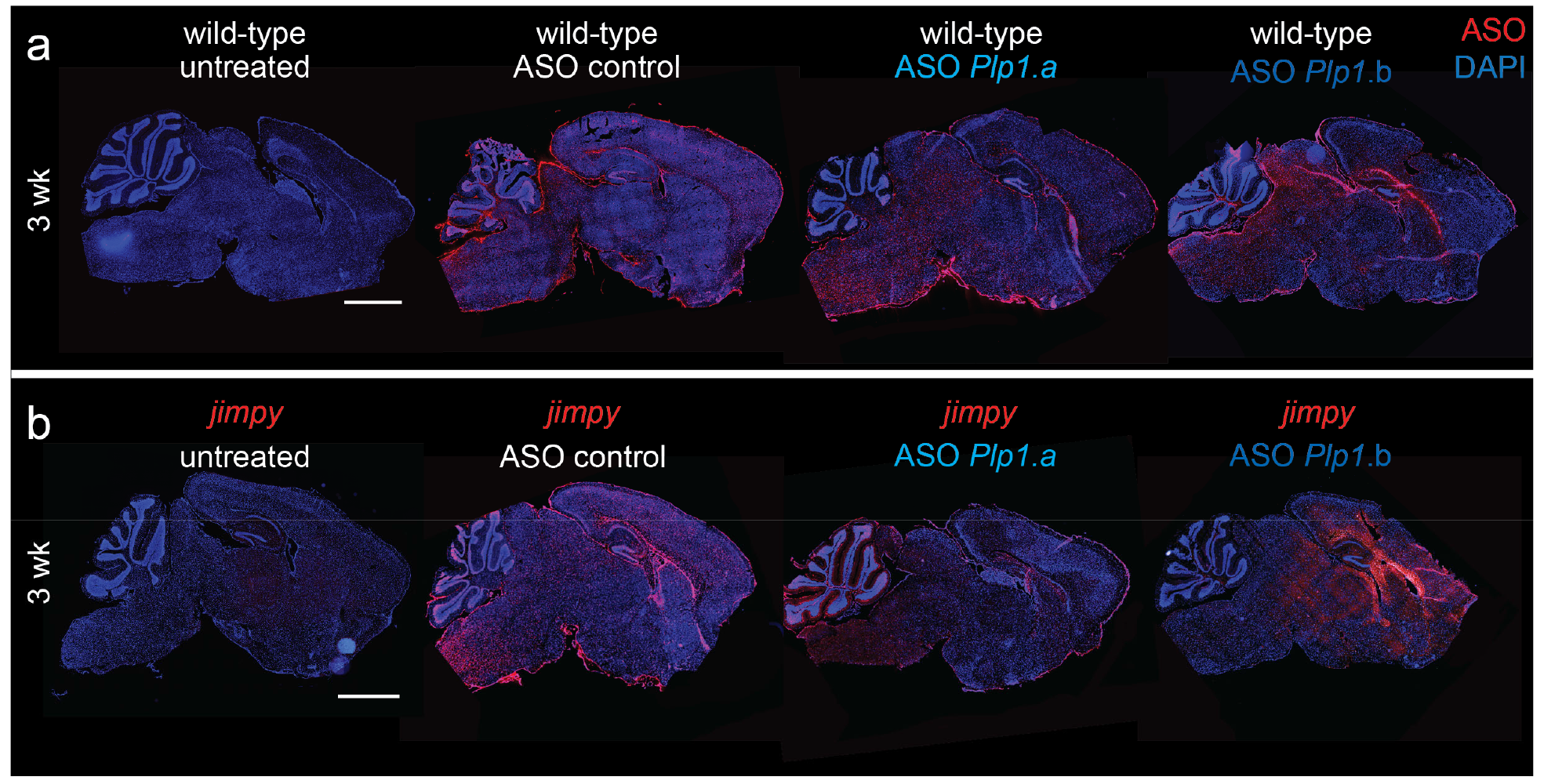
Plp1-targeted ASOs distribute widely throughout the CNS after ICV dosing in postnatal mice. **a-b**, Representative immunohistochemical images of 3 week whole-brain sagittal sections showing ASO (red) and total DAPI+ cells (blue) in wild-type mice (a) or *jimpy* (b) mice treated with ASO-*Plp1*.a, ASO-*Plp1*.b, or a control ASO by ICV injection at birth. Scale bar, 2mm.

**Extended Data Fig. 7.**
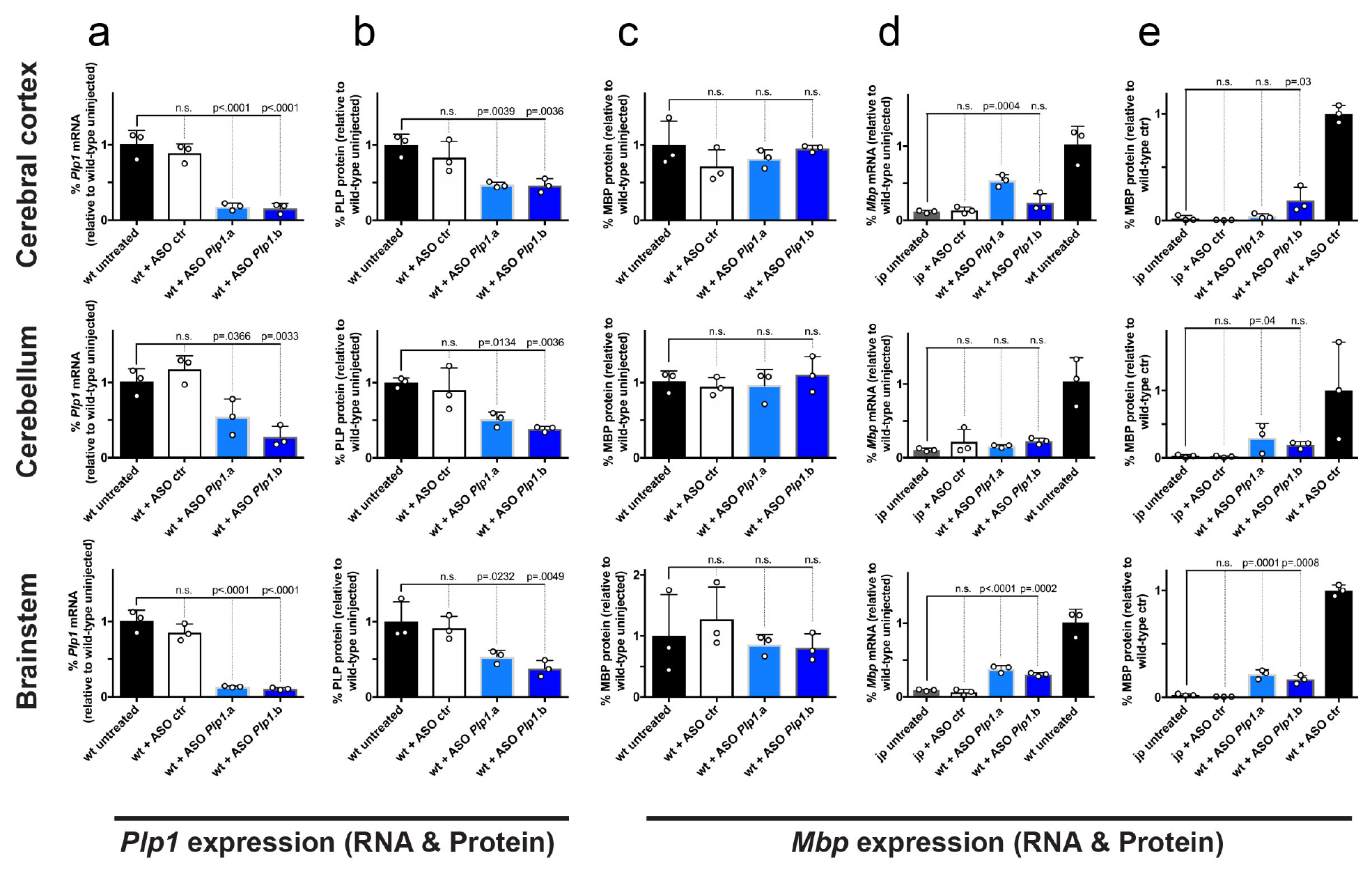
*Plp1*-targeted ASOs increase MBP transcript and protein levels in *jimpy* mice. **a**, RT-qPCR data showing the level of *Plp1* transcript suppression in 3 week old cerebral cortex, cerebellum, and brainstem of wild-type mice treated with the indicated ASOs (30ug dose) or controls at birth. Primer sites span *Plp1* exons 2-3, upstream of the *jimpy* mutation site. Individual data points represent the mean value of 4 technical replicates for each biological replicate (n=3 separate mice). Error bars show mean ± standard deviation. p-values calculated using one-way ANOVA with Dunnett’s correction for multiple comparisons. **b**, Western blot data showing the levels of PLP in 3 week old cerebral cortex, cerebellum, and brainstem of wild-type mice treated with the indicated ASOs (30ug dose) or controls at birth. Individual data points represent biological replicates (n=3 separate mice; see Supplementary Fig. 4 for full western blot images for all samples). Error bars show mean ± standard deviation. p-values calculated using one-way ANOVA with Dunnett’s correction for multiple comparisons. **c**, Western blot data showing the levels of MBP in 3 week old cerebral cortex, cerebellum, and brainstem of wild-type mice treated with the indicated ASOs (30ug dose) or controls at birth. Individual data points represent biological replicates (n=3 separate mice; see Supplementary Fig. 4 for full western blot images for all samples). Error bars show mean ± standard deviation. p-values calculated using one-way ANOVA with Dunnett’s correction for multiple comparisons. **d**, RT-qPCR data showing the level of *Mbp* transcript in 3 week old cerebral cortex, cerebellum, and brainstem of *jimpy* mice treated with the indicated ASOs (30ug dose) or controls at birth. Individual data points represent the mean value of 4 technical replicates for each biological replicate (n=3 separate mice). Error bars show mean ± standard deviation. p-values calculated using one-way ANOVA with Dunnett’s correction for multiple comparisons. **e**, Western blot data showing the level of MBP in 3 week old cerebral cortex, cerebellum, and brainstem of *jimpy* mice treated with the indicated ASOs (30ug dose) or controls at birth. Individual data points represent biological replicates (n=3 separate mice; see Supplementary Fig. 4 for full western blot images for all samples). Error bars show mean ± standard deviation. p-values calculated using one-way ANOVA with Dunnett’s correction for multiple comparisons.

**Extended Data Table 1.** Table containing details of published reports of *PLP1*-null patients.

| **PLP1 mutation** | **Patient #** | **Motor function** | **Cognitive function** | **Death** | **Ref.** |
| --- | --- | --- | --- | --- | --- |
| Frameshift and early termination codon (c.4delG)<br>*Protein absence confirmed by western blot* | 1 | Wheelchair bound at 35 with spasticity in all limbs | No intellectual disability and verbally communicated into mid-20s. Cooperative, alert, and aware of surroundings in adulthood. | Age 47 | (1, 2) |
|  | 2 | Wheelchair bound at 37 with spasticity in all limbs | No intellectual disability and verbally communicated into mid-20s. Cooperative, alert, and aware of surroundings in adulthood. | Age 49 |  |
|  | 3 | Walked from age 4-12, spasticity in all limbs at age 25 | Language delay. At 22 communicated, was cooperative, and alert. At 25 could follow commands, count, and distinguish colors. | Age 34 |  |
|  | 4 | Wheelchair bound at 17, gait deterioration starting at age 8, and spasticity in legs but not arms. | Able to read, follow commands, and perform arithmetic. | None reported |  |
| Frameshift and early termination codon c.191+1G>A<br>*No biochemical analysis* | 1 | Walked from age 6-9. Spastic with cerebellar dysfunction. | Attends normal school and can read and write. | None reported | (3) |
| Loss of start codon (c.3G>A)<br>*No biochemical analysis* | 1 | Early life motor dysfunction with spasticity in arms and legs. Further decline at age 33. | Early life intellectual disability. Further decline at age 33. | None reported | (2, 4) |
|  | 2 | Early life hypertonia and spasticity in arms and legs. Wheelchair bound at 6. | Language delay. Moderately retarded development. | None reported |  |
|  | 3 | Never walked and spasticity in legs by age 1, progressing to arms with age. | Mild cognitive deficits at age 20. | None reported | (5) |

**Supplementary Figure 1.**
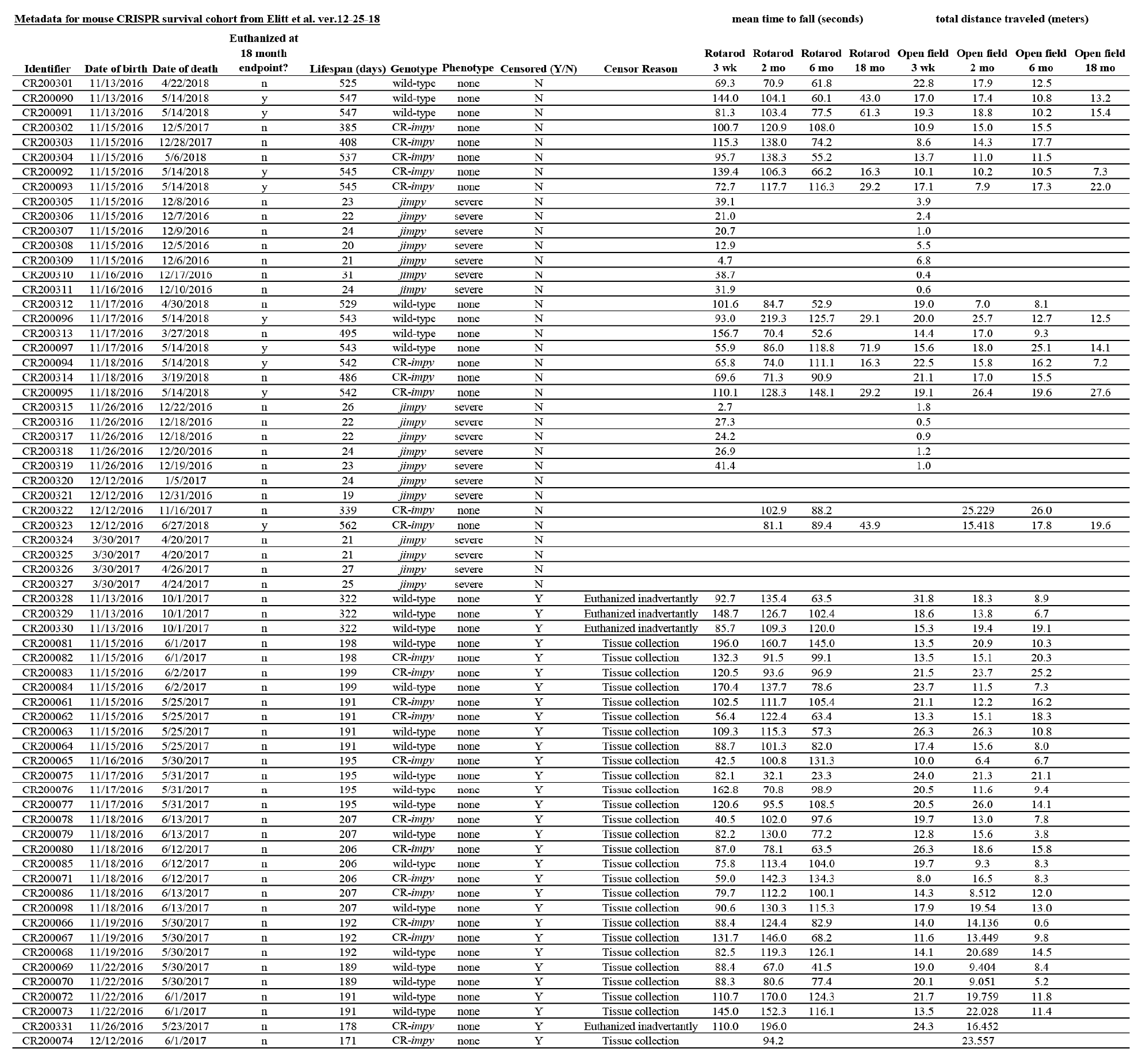
Table of metadata for all mice in Kaplan-Meier survival plot in Fig. 1b. Also included are raw data values for rotarod and open field assays in Fig. 2c, d.

**Supplementary Figure 2.**
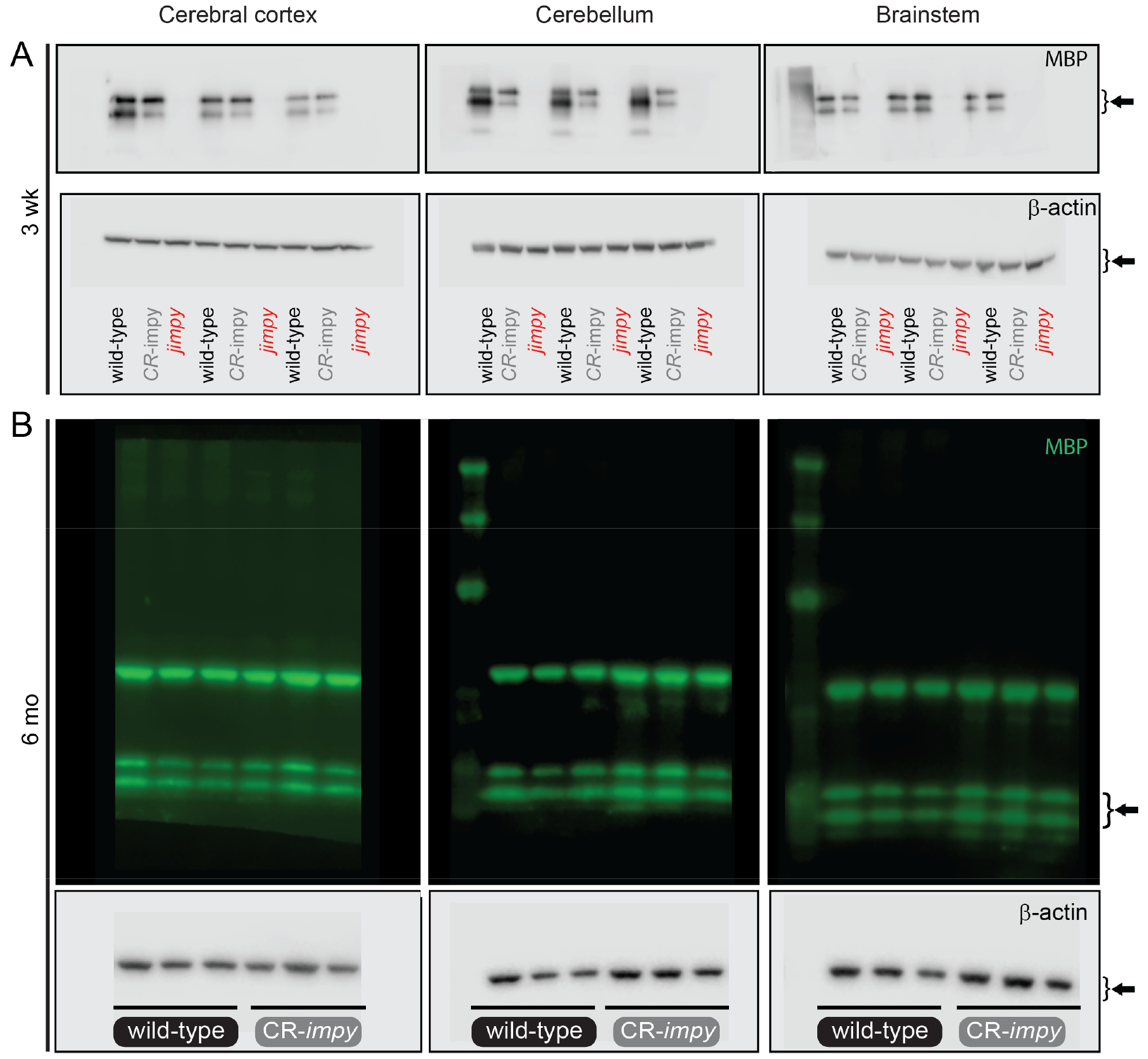
**a-b**, Labeled raw images of western blots for all samples in Extended Data Figs. 1d, f. The upper bands in the fluorescent blots in panel B are carry over from chemiluminescent detection of B-actin (bottom panel).

**Supplementary Figure 3.**
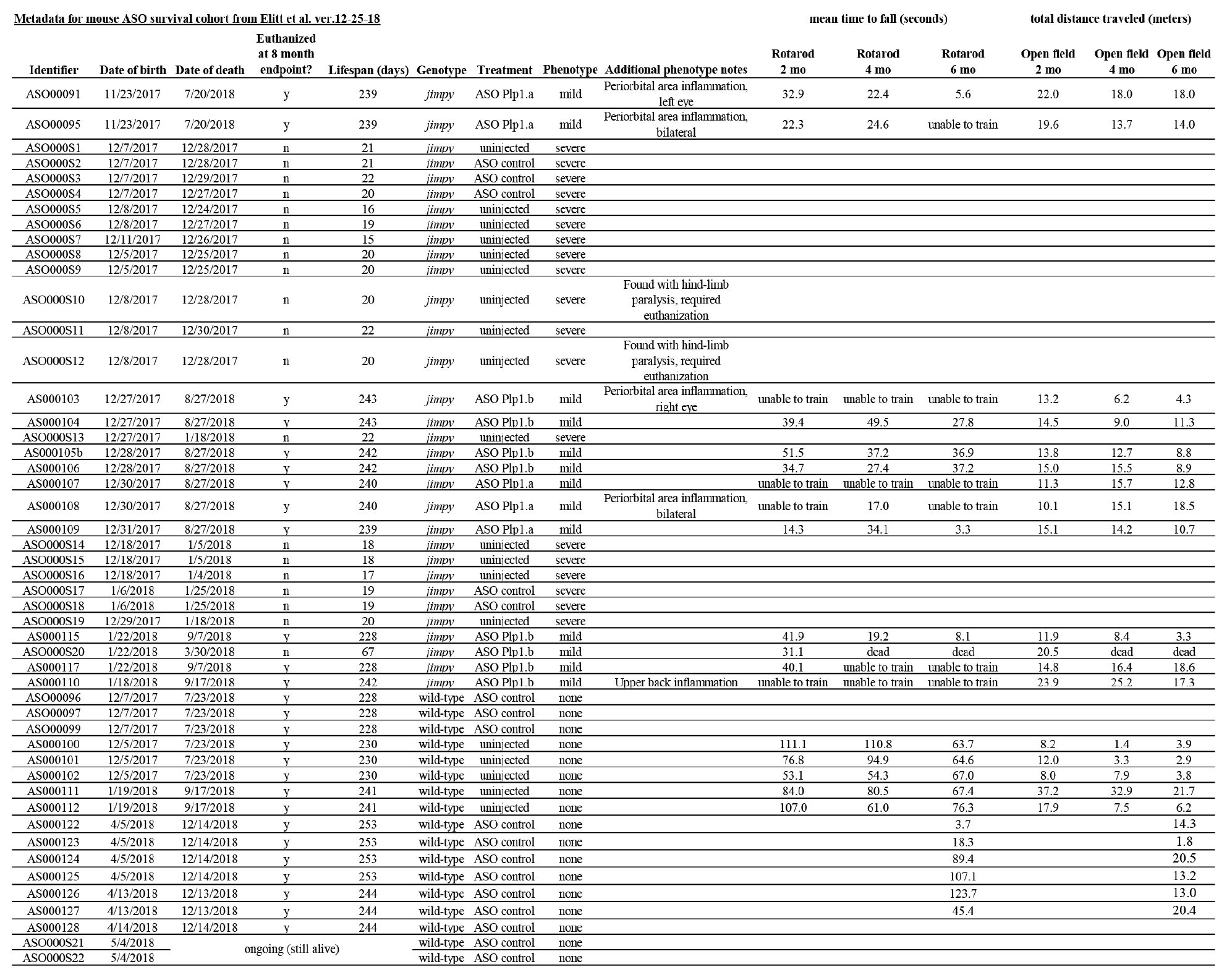
Table of metadata for all mice in Kaplan-Meier survival plot in Fig. 4c (as of 12-25-18 with two mice not yet at the 8 month endpoint). We noted 5 of 13 of ASO-treated *jimpy* mice in our survival cohort developed periorbital or upper back skin inflammation. The underlying cause of this is unknown as it was not observed in ASO treated wild-type littermates. Also included are raw data values for rotarod and open field assays in Fig. 4i, j.

**Supplementary Figure 4.**
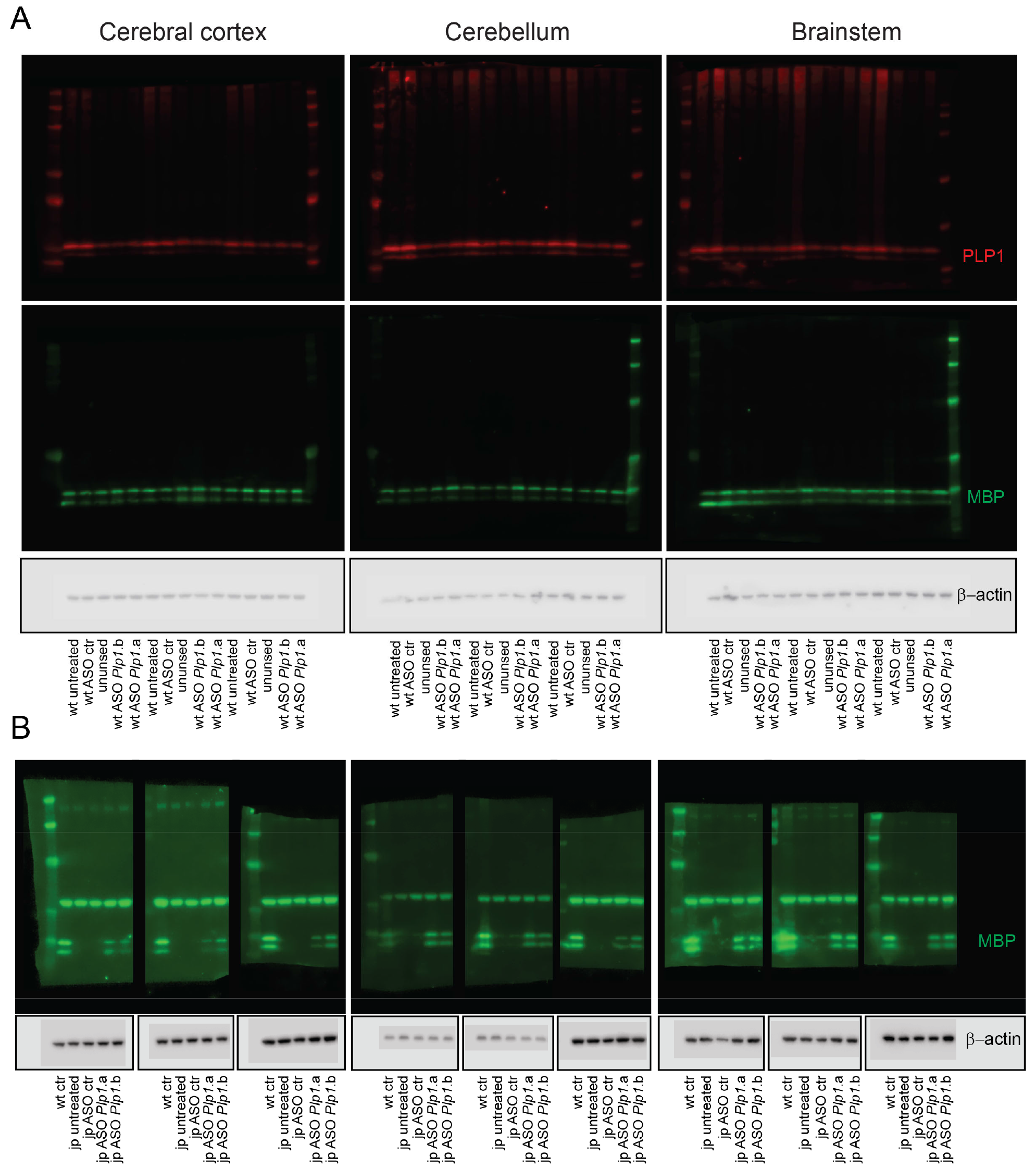
**a-b**, Labeled raw images of western blots for all samples in Extended Data Figs. 7b, c, e. The upper bands in the fluorescent blots in panel B are carry over from chemiluminescent detection of B-actin (bottom panel).

## SUPPLEMENTARY VIDEOS 1-4 (four separate MP4 files)

**Supplementary Video 1 | CRISPR-mediated knockdown of *Plp1* in *jimpy* mice rescues neurological phenotypes at 3 weeks of age**.

Video comparison of wild-type, *jimpy*, and CR-*impy* mice at 3 weeks of age.

**Supplementary Video 2 | CRISPR-mediated knockdown of *Plp1* in *jimpy* mice shows sustained rescue of neurological phenotypes at 18 months of age**.

Video comparison of wild-type and CR-*impy* mice at 18 weeks of age (study endpoint).

**Supplementary Video 3 | Postnatal delivery of **Plp1*-targeted* ASOs to *jimpy* mice rescues neurological phenotypes at 3 weeks of age.**

Video comparison of wild-type and *jimpy* mice treated with control and *Plp1*-targeting ASOs at 3 weeks of age.

**Supplementary Video 4 | Postnatal delivery of *Plp1*-targeted ASOs to *jimpy* mice shows sustained rescue of neurological phenotypes at 6 months of age.**

Video comparison of wild-type and *jimpy* mice treated with control and *Plp1*-targeting ASOs at 6 months of age.

## References

1 Goldman, S. A., Nedergaard, M. & Windrem, M. S. Glial progenitor cell-based treatment and modeling of neurological disease. Science 338, 491–495, doi:10.1126/science.1218071 (2012).

2 Gupta, N. et al. Neural stem cell engraftment and myelination in the human brain. Sci Transl Med 4, 155ra137, doi:10.1126/scitranslmed.3004373 (2012).

3 Saher, G. et al. Therapy of Pelizaeus-Merzbacher disease in mice by feeding a cholesterol-enriched diet. Nat Med 18, 1130–1135, doi:10.1038/nm.2833 (2012).

4 Prukop, T. et al. Progesterone antagonist therapy in a Pelizaeus-Merzbacher mouse model. Am JHum Genet 94, 533–546, doi:10.1016/j.ajhg.2014.03.001 (2014).

5 Wishnew, J. et al. Umbilical cord blood transplantation to treat Pelizaeus-Merzbacher Disease in 2 young boys. Pediatrics 134, e1451–1457, doi:10.1542/peds.2013-3604 (2014).

6 Tantzer, S., Sperle, K., Kenaley, K., Taube, J. & Hobson, G. M. Morpholino Antisense Oligomers as a Potential Therapeutic Option for the Correction of Alternative Splicing in PMD, SPG2, and HEMS. Mol Ther Nucleic Acids 12, 420–432, doi:10.1016/j.omtn.2018.05.019 (2018).

7 Cailloux, F. et al. Genotype-phenotype correlation in inherited brain myelination defects due to proteolipid protein gene mutations. Clinical European Network on Brain Dysmyelinating Disease. Eur J Hum Genet 8, 837–845, doi:10.1038/sj.ejhg.5200537 (2000).

8 Inoue, K. PLP1-related inherited dysmyelinating disorders: Pelizaeus-Merzbacher disease and spastic paraplegia type 2. Neurogenetics 6, 1–16, doi:10.1007/s10048-004-0207-y (2005).

9 Hobson, G. M. & Garbern, J. Y. Pelizaeus-Merzbacher disease, Pelizaeus-Merzbacher-like disease 1, and related hypomyelinating disorders. Semin Neurol 32, 62–67, doi:10.1055/s-0032-1306388 (2012).

10 Nave, K. A., Lai, C., Bloom, F. E. & Milner, R. J. Splice site selection in the proteolipid protein (PLP) gene transcript and primary structure of the DM-20 protein of central nervous system myelin. Proceedings of the National Academy of Sciences of the United States of America 84, 5665–5669 (1987).

11 Garbern, J. Y. Pelizaeus-Merzbacher disease: Genetic and cellular pathogenesis. Cell Mol Life Sci 64, 50–65, doi:10.1007/s00018-006-6182-8 (2007).

12 Baumann, N. & Pham-Dinh, D. Biology of oligodendrocyte and myelin in the mammalian central nervous system. Physiol Rev 81, 871–927 (2001).

13 Griffiths, I. et al. Axonal swellings and degeneration in mice lacking the major proteolipid of myelin. Science 280, 1610–1613 (1998).

14 Klugmann, M. et al. Assembly of CNS myelin in the absence of proteolipid protein. Neuron 18, 59–70 (1997).

15 Garbern, J. Y. et al. Patients lacking the major CNS myelin protein, proteolipid protein 1, develop length-dependent axonal degeneration in the absence of demyelination and inflammation. Brain 125, 551–561 (2002).

16 Hobson, G. M. & Kamholz, J. in GeneReviews((R)) (eds M. P. Adam et al.) (1993).

17 Jinek, M. et al. A programmable dual-RNA-guided DNA endonuclease in adaptive bacterial immunity. Science 337, 816–821, doi:10.1126/science.1225829 (2012).

18 Cong, L. et al. Multiplex genome engineering using CRISPR/Cas systems. Science 339, 819–823, doi:10.1126/science.1231143 (2013).

19 Dhaunchak, A. S. & Nave, K. A. A common mechanism of PLP/DM20 misfolding causes cysteine-mediated endoplasmic reticulum retention in oligodendrocytes and Pelizaeus-Merzbacher disease. Proceedings of the National Academy of Sciences of the United States of America 104, 17813–17818, doi:10.1073/pnas.0704975104 (2007).

20 Gow, A., Southwood, C. M. & Lazzarini, R. A. Disrupted proteolipid protein trafficking results in oligodendrocyte apoptosis in an animal model of Pelizaeus-Merzbacher disease. J Cell Biol 140, 925–934 (1998).

21 Nevin, Z. S. et al. Modeling the Mutational and Phenotypic Landscapes of Pelizaeus-Merzbacher Disease with Human iPSC-Derived Oligodendrocytes. Am J Hum Genet 100, 617–634, doi:10.1016/j.ajhg.2017.03.005 (2017).

22 Elitt, M. S. et al. Chemical Screening Identifies Enhancers of Mutant Oligodendrocyte Survival and Unmasks a Distinct Pathological Phase in Pelizaeus-Merzbacher Disease. Stem Cell Reports 11, 711–726, doi:10.1016/j.stemcr.2018.07.015 (2018).

23 Luders, K. A., Patzig, J., Simons, M., Nave, K. A. & Werner, H. B. Genetic dissection of oligodendroglial and neuronal Plp1 function in a novel mouse model of spastic paraplegia type 2. Glia 65, 1762–1776, doi:10.1002/glia.23193 (2017).

24 Miller, M. J., Kangas, C. D. & Macklin, W. B. Neuronal expression of the proteolipid protein gene in the medulla of the mouse. JNeurosci Res 87, 2842–2853, doi:10.1002/jnr.22121 (2009).

25 Lager, A. M. et al. Rapid functional genetics of the oligodendrocyte lineage using pluripotent stem cells. Nat Commun 9, 3708, doi:10.1038/s41467-018-06102-7 (2018).

26 Najm, F. J. et al. Rapid and robust generation of functional oligodendrocyte progenitor cells from epiblast stem cells. Nature methods 8, 957–962, doi:10.1038/nmeth.1712 (2011).

27 Lee, B. et al. Nanoparticle delivery of CRISPR into the brain rescues a mouse model of fragile X syndrome from exaggerated repetitive behaviours. Nature Biomedical Engineering, doi:10.1038/s41551-018-0252-8 (2018).

28 Anderson, K. R. et al. CRISPR off-target analysis in genetically engineered rats and mice. Nature methods 15, 512–514, doi:10.1038/s41592-018-0011-5 (2018).

29 Tsai, S. Q. & Joung, J. K. Defining and improving the genome-wide specificities of CRISPR-Cas9 nucleases. Nature reviews. Genetics 17, 300–312, doi:10.1038/nrg.2016.28 (2016).

30 Kosicki, M., Tomberg, K. & Bradley, A. Repair of double-strand breaks induced by CRISPR-Cas9 leads to large deletions and complex rearrangements. Nature biotechnology 36, 765–771, doi:10.1038/nbt.4192 (2018).

31 van Overbeek, M. et al. DNA Repair Profiling Reveals Nonrandom Outcomes at Cas9-Mediated Breaks. Molecular cell 63, 633–646, doi:10.1016/j.molcel.2016.06.037 (2016).

32 Rinaldi, C. & Wood, M. J. A. Antisense oligonucleotides: the next frontier for treatment of neurological disorders. Nat Rev Neurol 14, 9–21, doi:10.1038/nrneurol.2017.148 (2018).

33 Bennett, C. F., Baker, B. F., Pham, N., Swayze, E. & Geary, R. S. Pharmacology of Antisense Drugs. Annu Rev Pharmacol Toxicol 57, 81–105, doi:10.1146/annurev-pharmtox-010716-104846 (2017).

34 Finkel, R. S. et al. Treatment of infantile-onset spinal muscular atrophy with nusinersen: a phase 2, open-label, dose-escalation study. Lancet 388, 3017–3026, doi:10.1016/S0140-6736(16)31408-8 (2016).

35 Hagemann, T. L. et al. Antisense suppression of glial fibrillary acidic protein as a treatment for Alexander disease. Ann Neurol 83, 27–39, doi:10.1002/ana.25118 (2018).

36 Finkel, R. S. et al. Nusinersen versus Sham Control in Infantile-Onset Spinal Muscular Atrophy. N Engl JMed 377, 1723–1732, doi:10.1056/NEJMoa1702752 (2017).

37 Kordasiewicz, H. B. et al. Sustained therapeutic reversal of Huntington’s disease by transient repression of huntingtin synthesis. Neuron 74, 1031–1044, doi:10.1016/j.neuron.2012.05.009 (2012).

38 Crooke, S. T., Witztum, J. L., Bennett, C. F. & Baker, B. F. RNA-Targeted Therapeutics. CellMetab 27, 714–739, doi:10.1016/j.cmet.2018.03.004 (2018).

39 DeVos, S. L. et al. Tau reduction prevents neuronal loss and reverses pathological tau deposition and seeding in mice with tauopathy. Sci TranslMed 9, doi:10.1126/scitranslmed.aag0481 (2017).

40 Miller, T. M. et al. An antisense oligonucleotide against SOD1 delivered intrathecally for patients with SOD1 familial amyotrophic lateral sclerosis: a phase 1, randomised, first-inman study. Lancet Neurol 12, 435–442, doi:10.1016/S1474-4422(13)70061-9 (2013).

41 Becker, L. A. et al. Therapeutic reduction of ataxin-2 extends lifespan and reduces pathology in TDP-43 mice. Nature 544, 367–371, doi:10.1038/nature22038 (2017).

42 Meng, L. et al. Towards a therapy for Angelman syndrome by targeting a long noncoding RNA. Nature 518, 409–412, doi:10.1038/nature13975 (2015).

43 Passini, M. A. et al. Antisense oligonucleotides delivered to the mouse CNS ameliorate symptoms of severe spinal muscular atrophy. Sci Transl Med 3, 72ra18, doi:10.1126/scitranslmed.3001777 (2011).

44 Scoles, D. R. et al. Antisense oligonucleotide therapy for spinocerebellar ataxia type 2. Nature 544, 362–366, doi:10.1038/nature22044 (2017).

45 Gould, E. A. et al. Mild myelin disruption elicits early alteration in behavior and proliferation in the subventricular zone. eLife 7, doi:10.7554/eLife.34783 (2018).

46 Hsu, P. D. et al. DNA targeting specificity of RNA-guided Cas9 nucleases. Nature biotechnology 31, 827–832, doi:10.1038/nbt.2647 (2013).

47 Nakagata, N., Okamoto, M., Ueda, O. & Suzuki, H. Positive effect of partial zona-pellucida dissection on the in vitro fertilizing capacity of cryopreserved C57BL/6J transgenic mouse spermatozoa of low motility. Biol Reprod 57, 1050–1055 (1997).

48 Schmid-Burgk, J. L. et al. OutKnocker: a web tool for rapid and simple genotyping of designer nuclease edited cell lines. Genome Res 24, 1719–1723, doi:10.1101/gr.176701.114 (2014).

49 Glascock, J. J. et al. Delivery of therapeutic agents through intracerebroventricular (ICV) and intravenous (IV) injection in mice. J Vis Exp, doi:10.3791/2968 (2011).

## Supplementary References

1. J. Y. Garbern et al., Proteolipid protein is necessary in peripheral as well as central myelin. Neuron 19, 205–218 (1997).

2. J. Y. Garbern et al., Patients lacking the major CNS myelin protein, proteolipid protein 1, develop length-dependent axonal degeneration in the absence of demyelination and inflammation. Brain 125, 551–561 (2002).

3. P. Lassuthova et al., Three new PLP1 splicing mutations demonstrate pathogenic and phenotypic diversity of Pelizaeus-Merzbacher disease. Journal of child neurology 29, 924–931 (2014).

4. E. A. Sistermans et al., A (G-to-A) mutation in the initiation codon of the proteolipid protein gene causing a relatively mild form of Pelizaeus-Merzbacher disease in a Dutch family. Hum Genet 97, 337–339 (1996).

5. C. K. Hand, G. Bernard, M.-P. Dubé, M. I. Shevell, G. A. Rouleau, A Novel PLP1 Mutation Further Expands the Clinical Heterogeneity at the Locus. The Canadian Journal of Neurological Sciences 39, 220–224 (2014).

